# Widespread epistasis among beneficial genetic variants revealed by high-throughput genome editing

**DOI:** 10.1101/2022.06.05.494911

**Authors:** Roy Moh Lik Ang, Shi-An Anderson Chen, Alexander F. Kern, Yihua Xie, Hunter B. Fraser

## Abstract

Genetic interactions occur when a variant’s phenotypic effect is altered by variation at other genomic loci. Also known as epistasis, these interactions shape the genetic architecture of complex traits and modify phenotypes across genetic backgrounds. However, the factors associated with their occurrence remain poorly understood. To investigate this, we employed high-throughput genome editing to measure the fitness effects of 1,826 naturally polymorphic variants in four genetically diverse strains of *Saccharomyces cerevisiae*. About 31% of variants affect fitness in a common laboratory environment, of which 24% have strain-specific fitness effects indicative of epistasis. We found that beneficial variants are more likely to exhibit genetic interactions, and that genetic interactions are depleted among variants at higher allele frequencies. In addition, we demonstrate that these epistatic interactions for fitness can be mediated by specific traits such as flocculation ability. This work suggests that adaptive evolution from standing variation will often involve trade-offs where a variant is only beneficial in some genetic backgrounds, potentially explaining why many beneficial variants remain polymorphic. In sum, we provide a framework to understand the factors influencing epistasis in natural genetic variants with single-nucleotide resolution, revealing widespread epistasis among beneficial variants.

## Introduction

Genetic interactions arise when a mutation’s contribution to a trait is altered by variation at one or more other genomic loci. They are pervasive in the genetic architecture of complex traits, and collectively shape the genetic background’s impact on phenotype (Chandler et al., 2013; Cooper et al., 2013; Goldstein and Ehrenreich, 2021; Huang et al., 2012; Nadeau, 2001; Taylor and Ehrenreich, 2015). Dissecting the causes of genetic background effects is pivotal to understanding how it affects phenotypic variation within populations (Bergman and Siegal, 2003; Masel, 2013; Rutherford and Lindquist, 1998), evolutionary trajectories of deleterious and beneficial variants (Hemani et al., 2013; Johnson et al., 2019; Kryazhimskiy et al., 2014), as well as prediction and treatment of diseases with variable penetrance across individuals (Chen et al., 2016; Narasimhan et al., 2016; Riordan and Nadeau, 2017). However, the molecular mechanisms underlying genetic interactions remain poorly understood. Determining the genomic features associated with the incidence of genetic interactions is of particular interest, as it will facilitate the prediction of epistasis genome-wide based on individual genotypes.

The conventional approach to studying genetic interactions involves shuffling variation between two or more populations via recombination and mapping quantitative trait loci (QTL) in hundreds of offspring (Bloom et al., 2015; Ehrenreich, 2017; Huang et al., 2012). However, QTLs typically have insufficient resolution to identify the specific causal variants associated with the phenotype of interest, due to limitations in available recombination events (Rockman, 2012; Schaid et al., 2018). In addition, because QTL effect sizes are measured over many offspring, the nuances of genetic background effects are lost, which may lead to underestimation of the contribution of genetic interactions to phenotypic variation.

A more direct way to measure genetic interactions is to screen mutations in distinct genetic backgrounds (Dowell et al., 2010; Galardini et al., 2019). By introducing mutations into individuals with sufficient sequence divergence and measuring phenotypic effects in each genetic background, causal variants with background-specific effects can be identified. This is especially relevant in complex polygenic traits, where genetic background is expected to contribute more to the genotype-phenotype relationship than in highly penetrant monogenic phenotypes. Related approaches involve measuring genetic interactions between large collections of mutations (Costanzo et al., 2016; Kuzmin et al., 2018), or within single genes (Li et al., 2016; Puchta et al., 2016; Weinreich et al., 2006). However, it is impractical to screen all possible combinations of mutations at present, making it challenging to address the global epistasis question using a ground-up approach. Furthermore, most genetic screens focus on whole-gene knockouts or random mutations that produce large phenotypic effects, which are not representative of natural variation, and thus cannot reveal the extent to which epistasis shapes phenotypes in natural populations.

In this study, we seek to address this by screening 1,826 natural variants in four genetically diverse strains of *Saccharomyces cerevisiae* isolated from different ecological origins for effects on growth in a common laboratory environment. This was done using CRISPEY-BAR (Chen et al., submitted) – an improved high-throughput precision editing and lineage tracking platform based on the CRISPEY technology previously developed by Sharon et al. (2018). We detected genetic interactions in 24% of variants that affect fitness, and identified factors associated with their occurrence, laying the foundation for understanding the molecular underpinnings of epistasis across the genome.

## Results

### CRISPEY-BAR fitness screen of natural variants in four yeast strains

To quantitatively assess the extent to which genetic interactions influence complex traits, we screened 1,826 natural variants (Peter et al., 2018) in four genetically diverse yeast strains for effects on growth in a synthetic laboratory medium. We selected variants in and around 103 “hub” genes which were implicated in a high number of genetic interactions based on a previous genome-wide deletion screen (Costanzo et al., 2016). We cloned 5,319 guide-donor oligomers (Table S1) with unique barcodes into the CRISPEY-BAR vector, which executes CRISPEY precision editing of the target variant (Sharon et al., 2018) while simultaneously inserting a barcode at a predefined genomic locus (Levy et al., 2015) to track the edit within individual cells (Chen et al., submitted). The plasmid pool was independently transformed and edited in each yeast strain, and edited yeast pools were competed in synthetic complete media with 2% glucose for ∼60 generations. Barcodes were sampled every ∼15 generations to track the relative abundance of variants, from which the fitness effects can be estimated (Figure 1A).

**Figure 1.**
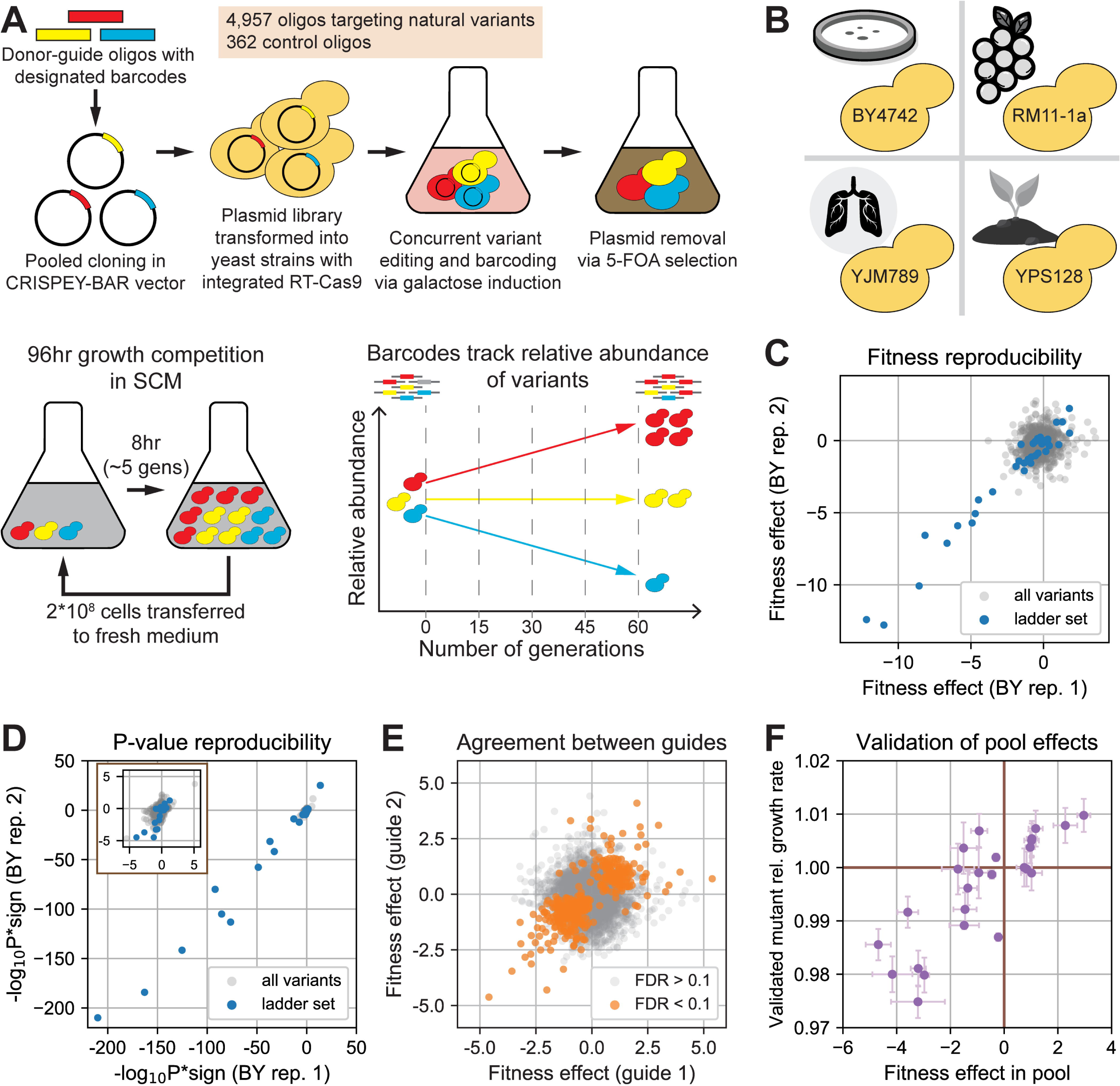
CRISPEY-BAR fitness screen of natural variants in four genetically distinct yeast strains. (A) Experiment design of CRISPEY-BAR pooled editing and growth competition. (B) Yeast strains used in fitness screen, with graphical representation of the ecological niche from which each strain was isolated. Graphics from stockunlimited.com. (C) Variant fitness effects are correlated between replicate growth competitions of independently edited pools. (D) P-values of variant fitness estimates are correlated between replicate growth competition. The fitness effects of 42 deletion mutations with known growth effects in BY (“ladder set”, blue) are highly reproducible. Inset: zoom-in on P-values for non-ladder variants. (E) Fitness effects measured from independent guides targeting each variant, across all growth competitions. Natural variants with significant fitness effects (orange) exhibit stronger correlation between guides than those indistinguishable from non-editing controls (gray). (F) Validation of 25 significant fitness effects from the pool growth competitions in sequence-verified mutants. Brown lines indicate wild-type fitness.

We selected four yeast strains (BY4742, RM11-1a, YPS128, YJM789) isolated from different ecological niches to maximize phenotypic diversity. Unlike the common laboratory strain BY4742 (BY), RM11-1a (RM) was derived from a vineyard isolate (Brem et al., 2002), YJM789 (YJM) was derived from a strain found in the lungs of an AIDS patient (Tawfik et al., 1989; Wei et al., 2007), and YPS128 (YPS) was isolated from the soil beneath *Quercus alba* (Sniegowski et al., 2002) (Figure 1B). The non-BY strains possess 42,000-62,000 SNPs relative to the reference strain S288c, giving them an average sequence divergence of ∼0.5% (Peter et al., 2018).

Variant fitness effects from each growth competition were measured relative to non-editing barcodes that do not edit the genome apart from the barcode insertion, which represent the wild-type distribution of neutral fitness measurements in the pool. Fitness effects can be directly compared across competitions by calculating a Z-score based on the distribution of neutral fitness values from non-editing barcodes in each competition (Table S2, see Methods). Applying this approach to the 1,826 natural variants in our screen, we identified 564 (31%) variants with significant non-neutral fitness effects in at least one strain (FDR < 0.1). To estimate the reproducibility of our measurements, we designed three types of replications. The first was replication of gene deletion fitness effects across competitions. For this, we included 42 “ladder” control oligomers that introduce short frameshift deletions in genes with known growth effects in BY (Breslow et al., 2008). We found that both variant fitness effects (Pearson’s r = 0.98, p < 10^-25^ for ladder set; Figure 1C) and their associated p-values (Pearson’s r = 0.99, p < 10^-30^ for ladder set; Figure 1D) are highly reproducible across replicate growth competitions conducted on independently edited BY pools.

The second replication was to assess the agreement between oligomers editing the same variant. We designed at least two oligomers with unique guide sequences to target each variant in the library, and found that their fitness effects are significantly correlated among non-neutral variants (Pearson’s r = 0.74, p < 10^-77^ for variants with FDR < 0.1; Figure 1E). There is no correlation among apparently neutral variants (Pearson’s r = -0.03 for variants with FDR > 0.1; Figure 1E), as expected since any variation in the fitness effects of neutral variants is solely measurement error, which is not expected to correlate across replicates. As a third type of replication, we validated 25 significant fitness effects by generating sequence-verified mutants in each genetic background and competing them individually with their non-edited parents. We found a significant correlation between the relative growth rate of validated mutants and their fitness effects in the pooled competitions (Pearson’s r = 0.81, p < 10^-6^; Figure 1F). Taken together, our results show that the growth competition fitness effects across yeast strains are accurate and highly reproducible.

### Global differences in the distribution of fitness effects across yeast strains

Next, we sought to compare the distribution of variant fitness effects (DFE) in each yeast strain. We found that DFEs are significantly different across the four strains (Kruskal-Wallis p < 10^-44^; Figure 2A); there are significant shifts in DFE between all strain pairs except BY/RM (pairwise Mann-Whitney U adjusted p < 10^-4^ after Bonferroni correction). More specifically, the median fitness effect in YJM (median = 0.038) is slightly greater than in non-YJM strains (median = -0.078 to -0.176), and more variants have their most beneficial fitness effect in YJM than in other strains (chi-square p < 10^-63^). One possible explanation for this could be that YJM is less well adapted to the conditions of our competitions, though this was not reflected in a lower baseline growth rate for YJM (Figure S1).

**Figure 2.**
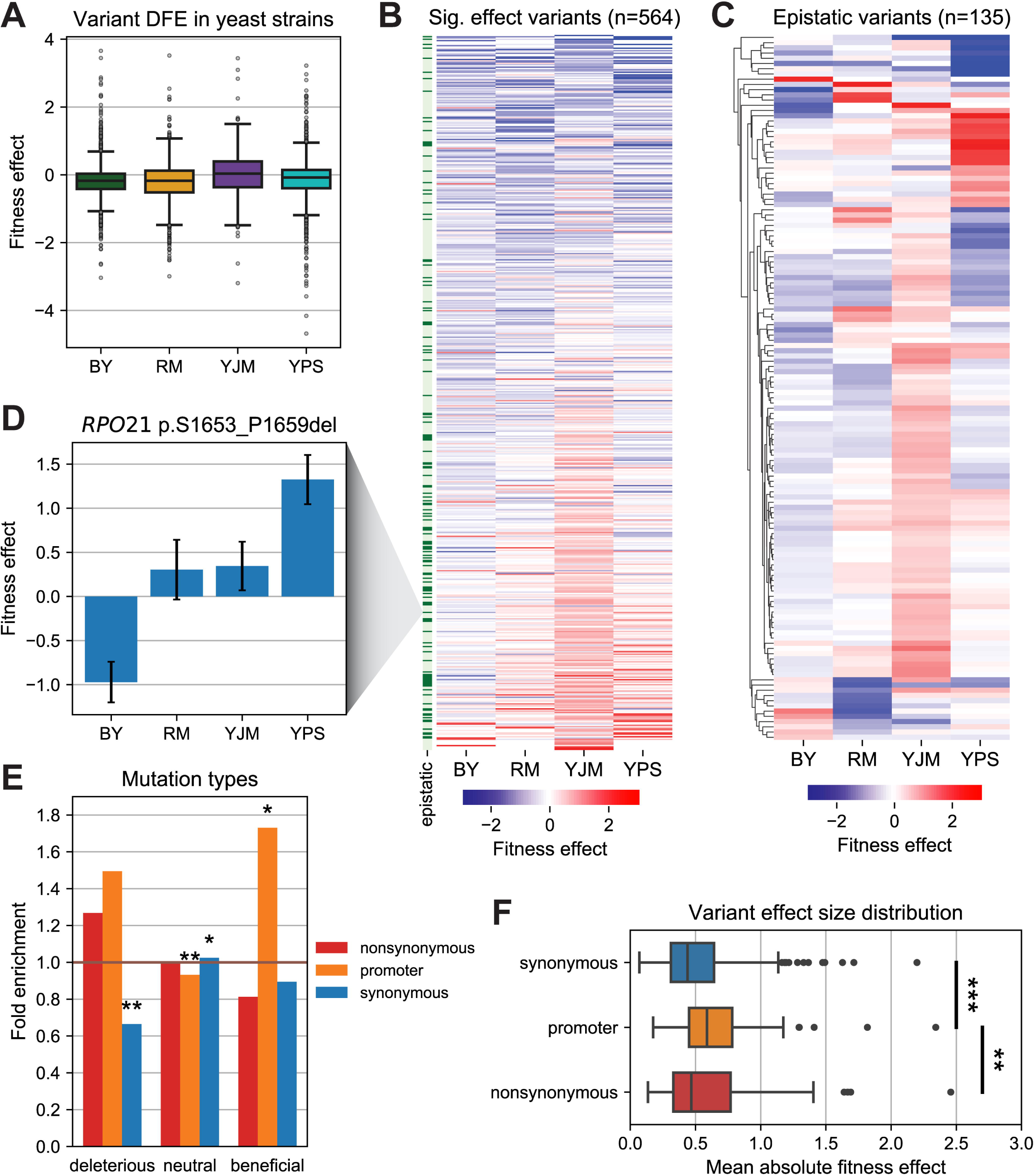
Global differences in the distribution of fitness effects across genetic backgrounds. (A) Distribution of fitness effects of 1,826 natural variants across four yeast strains. (B) Heatmap of fitness effects for 564 variants that are significantly different from neutral in at least one strain (FDR < 0.1), sorted by mean fitness effect. Row colors next to the heatmap indicate the presence (dark green) or absence (light green) of genetic interactions. (C) Heatmap of fitness effects for 135 epistatic variants with significant genetic interactions. (D) Example of sign epistasis between BY and YPS in an *RPO21* in-frame deletion mutation. Data is represented by mean ± SEM. (E) Deleterious variants (mean fitness effect < 5^th^ percentile) are depleted for synonymous variants, while beneficial variants (mean fitness effect > 95^th^ percentile) are enriched for promoter variants. Asterisks indicate significance levels for Fisher’s exact test. *p<0.05 **p<0.005 (F) The distribution of effect sizes among variants with significant fitness effects is significantly larger in promoter variants. Asterisks indicate significance levels for Mann-Whitney U test. **p<0.005 ***p<0.0005

Among variants with non-neutral fitness effects, we found substantial variability in effect size and direction between strains (Figure 2B), suggesting the widespread action of genetic background effects. To examine this more closely, we identified 135 epistatic variants with significant genetic interactions based on the differences in fitness effects between strains (F-test FDR < 0.05, and ≥ 2 pairwise fitness differences between strains at FDR < 0.1, Tables S3 and S4, see Methods), and observed multiple instances of strain-specific fitness effects (Figure 2C). These epistatic variants account for 7.9% of all variants measured in two or more strains.

However, among variants with significant fitness effects, the proportion of epistatic variants is 24%. Strikingly, 18% of the epistatic variants involve a significant change in the direction of fitness effect between at least two strains, a phenomenon known as sign epistasis (Weinreich et al., 2005). For example, we observed an in-frame deletion in a region of the C-terminal domain of RPO21 with an effect consistent with previously observed growth defects in BY (Babokhov et al., 2018) that produced a positive fitness effect in YPS, and had no significant effect in RM or YJM (Figure 2D).

We sorted variants by their mean fitness effect across all four strains to identify deleterious (mean fitness effect < 5^th^ percentile) and beneficial (mean fitness effect > 95^th^ percentile) variants in our library. We observed that synonymous variants are significantly depleted among deleterious variants (0.66-fold depletion, Fisher’s exact p = 0.0036) while promoter variants are significantly enriched among beneficial variants (1.73-fold enrichment, Fisher’s exact p = 0.007; Figure 2E). Nonsynonymous variants are also slightly enriched among deleterious variants, though it did not reach significance (1.27-fold enrichment, Fisher’s exact p = 0.06). In addition, we found that the distribution of four-strain mean absolute fitness effects for non-neutral variants is significantly larger in promoter variants than in synonymous variants (Mann-Whitney U p < 10^-4^), as well as nonsynonymous variants (Mann-Whitney U p = 0.0049; Figure 2F). In sum, our results reveal global differences in the distribution of fitness effects in natural variants across different genetic backgrounds, as well as between different types of mutations.

### Beneficial variants are more likely to exhibit genetic interactions

We next examined how genetic interactions relate to the variant mean fitness effects across the four strains. We found that the proportion of epistatic variants increases with greater mean fitness effect (Figures 3A and S2A). Although we would expect that genetic interactions are more readily detected in variants with larger effect sizes, we still find more genetic interactions among beneficial variants than deleterious variants with similar effect sizes (Figure S2B). This asymmetric distribution of epistasis becomes apparent when comparing the four-strain fitness landscape of deleterious variants (Figure 3B) to the fitness landscape of beneficial variants (Figure 3C): While deleterious effects tend to be shared across the yeast strains, beneficial effects are more often strain-specific, conferring a positive fitness effect in some strains while being neutral or slightly deleterious in others. Specifically, deleterious variants are 2.03-fold more likely to have concordant effects across strains than beneficial ones (Fisher’s exact p < 10^-5^). As a result, genetic interactions are 3.13-fold enriched in beneficial variants relative to deleterious variants (binomial p < 10^-6^) and 3.71-fold enriched relative to the library-wide average (Fisher’s exact p < 10^-8^; Figure 3D). Additionally, we found that epistatic variants were more likely to have their most beneficial and deleterious effects in specific genetic backgrounds. For instance, epistatic variants with negative mean fitness effect were 1.90-fold more likely to have their most deleterious effect in YPS (Fisher’s exact p = 0.00035). On the other hand, epistatic variants with positive mean fitness effect were 0.22-fold less likely to have their most beneficial effect in BY (Fisher’s exact p = 0.004; Figure S3).

**Figure 3.**
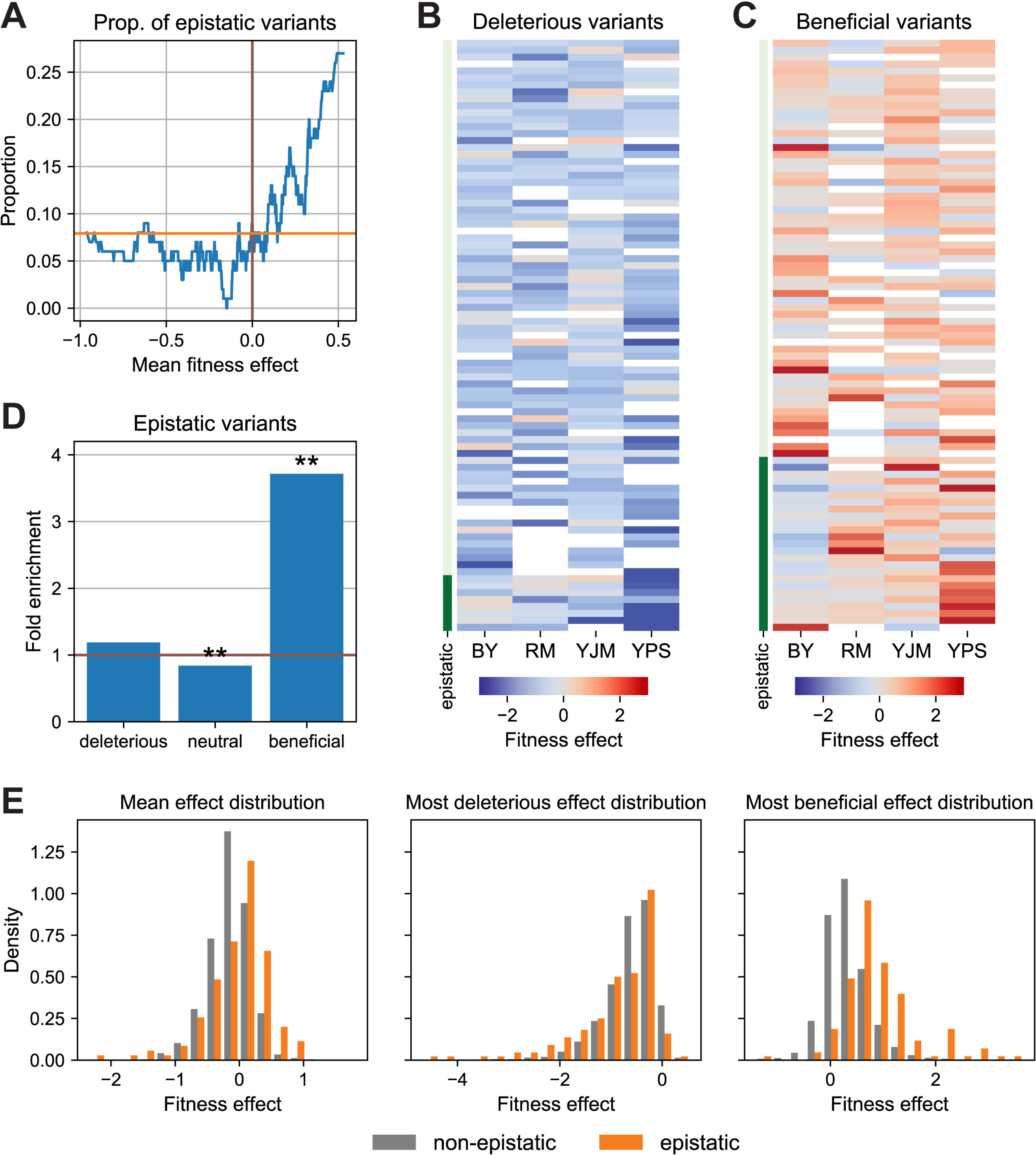
Beneficial variants are more likely to exhibit genetic interactions. (A) Proportion of variants with genetic interactions increases with greater mean fitness effect. Proportion is calculated from a 100-variant sliding window across all variants ranked by the mean fitness effect across four strains. Orange line indicates the library-wide mean proportion of epistatic variants. (B) Deleterious variants (mean fitness effect < 5^th^ percentile) show consistent negative effects across yeast strains. (C) Beneficial variants (mean fitness effect > 95^th^ percentile) are more likely to exhibit strain-specific fitness effects. Row colors next to the heatmap indicate the presence (dark green) or absence (light green) of genetic interactions. Missing values are colored white. (D) Genetic interactions are strongly enriched among beneficial variants. Asterisks indicate significance levels for Fisher’s exact test. *p<0.05 **p<0.005 (E) Distributions of mean effect and most beneficial effect are shifted positive for epistatic variants (orange) compared to non-epistatic (gray) variants, but not in the most deleterious effect distribution.

We ruled out several factors that may have caused the enrichment of genetic interactions among beneficial variants: For example, we found no significant difference in the number of missing fitness effect measurements between beneficial and deleterious variants (chi-square test p = 0.73) which could have created asymmetrical power to detect genetic interactions. In addition, positive effects from our fitness validation experiment are more strongly correlated with pool fitness measurements (Pearson’s r = 0.76, p = 0.018) than negative effects are (Pearson’s r = 0.68, p = 0.0038; Figure 1F), suggesting that experimental noise is not contributing to more spurious genetic interactions among beneficial variants. We also speculated that YJM’s slightly more positive DFE may be driving many strain-specific beneficial effects and inflating the number of epistatic variants with positive mean effect. However, the enrichment of genetic interactions among beneficial variants remains significant even when YJM is excluded from the results, which indicates that YJM is not the main driver behind the asymmetry in epistatic effects (Figure S4).

Consequently, we find significant shifts in the DFEs between epistatic and non-epistatic variants (Figure 3E), specifically positive shifts in the distribution of mean effect (Mann-Whitney U p < 10^-5^), as well as most beneficial effect (Mann-Whitney U p < 10^-30^), of variants across the four yeast strains. There is no significant difference in the distribution of most deleterious effect of variants, even after excluding neutral variants (Mann-Whitney U p = 0.78). Taken together, our results show the asymmetric distribution of epistasis along the distribution of fitness effects, with beneficial variants being more likely to exhibit genetic interactions.

### Genetic interactions are depleted at higher allele frequencies

If genetic interactions result in deleterious fitness effects in some strains, but beneficial effects in others, epistatic variants might exist at intermediate allele frequencies in the population. For example, a variant that is only beneficial in ∼10% of strains may be maintained by natural selection at ∼10% frequency, only being maintained in those strains where it is beneficial. Consistent with this, we observed enrichment for genetic interactions at allele frequencies between 5 to 10% (1.64-fold enrichment, Fisher’s exact p = 0.02; Figure 4A). In contrast, variants below 5% allele frequency are not enriched for genetic interactions, and those above 10% allele frequency are slightly depleted for genetic interactions compared to those at 5-10%; the allele frequency distribution is significantly different between epistatic and non-epistatic common variants (Kolmogorov-Smirnov test p < 0.05; Figure 4B).

**Figure 4.**
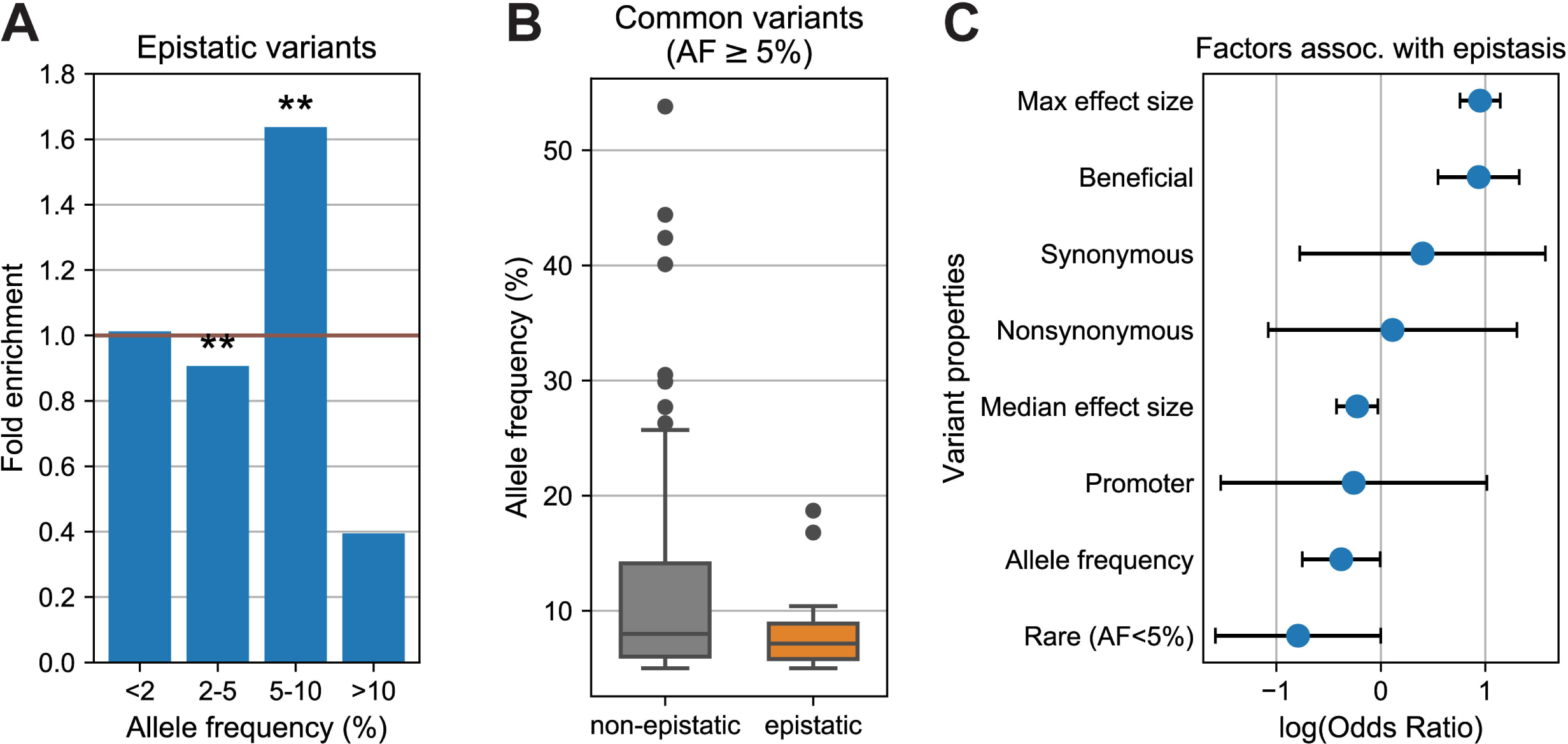
Genetic interactions are depleted at higher allele frequencies. (A) Variants with allele frequency (AF) of 5-10% are significantly enriched for genetic interactions. Allele frequencies were obtained from Peter et al. (2018). (B) Among common variants (AF ≥ 5%), the allele frequency distribution in epistatic variants is shifted lower relative to non-epistatic variants. (C) Multiple logistic regression odds ratios for variables associated with the incidence of genetic interactions in natural variants. Data is represented by mean log odds ratio ± 95% confidence interval from the logistic model.

Because many factors could potentially be associated with epistasis, we fitted a multiple logistic regression model with several covariates related to variant and gene properties to determine the association of each variable with the incidence of genetic interactions. We found that in addition to a variant’s median and maximum effect size, its median effect direction and allele frequency are significant predictors for epistasis in our data: Specifically, variants that are beneficial and have a lower allele frequency are more likely to exhibit genetic interactions (Figure 4C). Properties of the gene associated with each variant such as the number of genetic interactions the gene is involved in (Costanzo et al., 2016), its essentiality in S288c, and its paralog status have no significant association with epistasis (Figure S5).

Due to the uneven sample sizes of the 1,011 yeast strains across ecological origins, we wondered if population structure might be associated with the incidence of genetic interactions among natural variants; if epistatic variants are typically beneficial in one genetic background, but not others, we would expect epistatic variants to be concentrated in fewer ecological origins than non-epistatic variants. However, we found no significant difference in the number of ecological origins represented between epistatic and non-epistatic variants (Figure S6). To examine whether variants in specific ecological niches were more likely to exhibit genetic interactions, we ran a principal components analysis (PCA) on the frequency at which a variant appears in each ecological origin of the 1,011 yeast strains collection (Peter et al., 2018) and fitted the first six principal components accounting for 74.8% of variance in the PCA as covariates in the logistic model (Figure S7). Only PC1, explaining 41.6% of variance in the PCA, had a significant association with epistasis (Figure S8A), and we found that PC1 was highly correlated with allele frequency (Pearson’s r = 0.96, p < 10^-100^; Figure S8B), suggesting that in terms of population structure, allele frequency alone is the major determinant of the distribution of genetic interactions across natural variants.

### Genetic interactions are concentrated in genes associated with physiological differences between yeast strains

Genetic variation affects phenotype through the modification of gene function, e.g. through protein sequence alterations or gene expression changes. Hence, examining the distribution of genetic interactions among genes might reveal key distinctions in cellular physiology across the four yeast strains. We quantified the epistasis signal – defined as the extent of the difference in fitness effect across four strains based on the F-test p-value – for variants with significant fitness effects and found that genetic interactions are not randomly distributed across genes, but instead are concentrated in a few key genes (Kruskal-Wallis p = 0.00042; Figure 5A). More specifically, variants in a gene that suppresses flocculation (*SFL1*) showed significantly greater epistasis signal than the rest of the library (Mann-Whitney U p = 0.00049). Genetic interactions were observed in both coding and noncoding variants of *SFL1*, with several missense variants exhibiting strong epistasis across the four strains (Figure 5B). The high density of variants with strong genetic interactions in *SFL1* is contrasted by other genes in our study such as *POL2*, which show little to no epistasis signal despite having multiple variants with significant fitness effects (Figure 5C).

**Figure 5.**
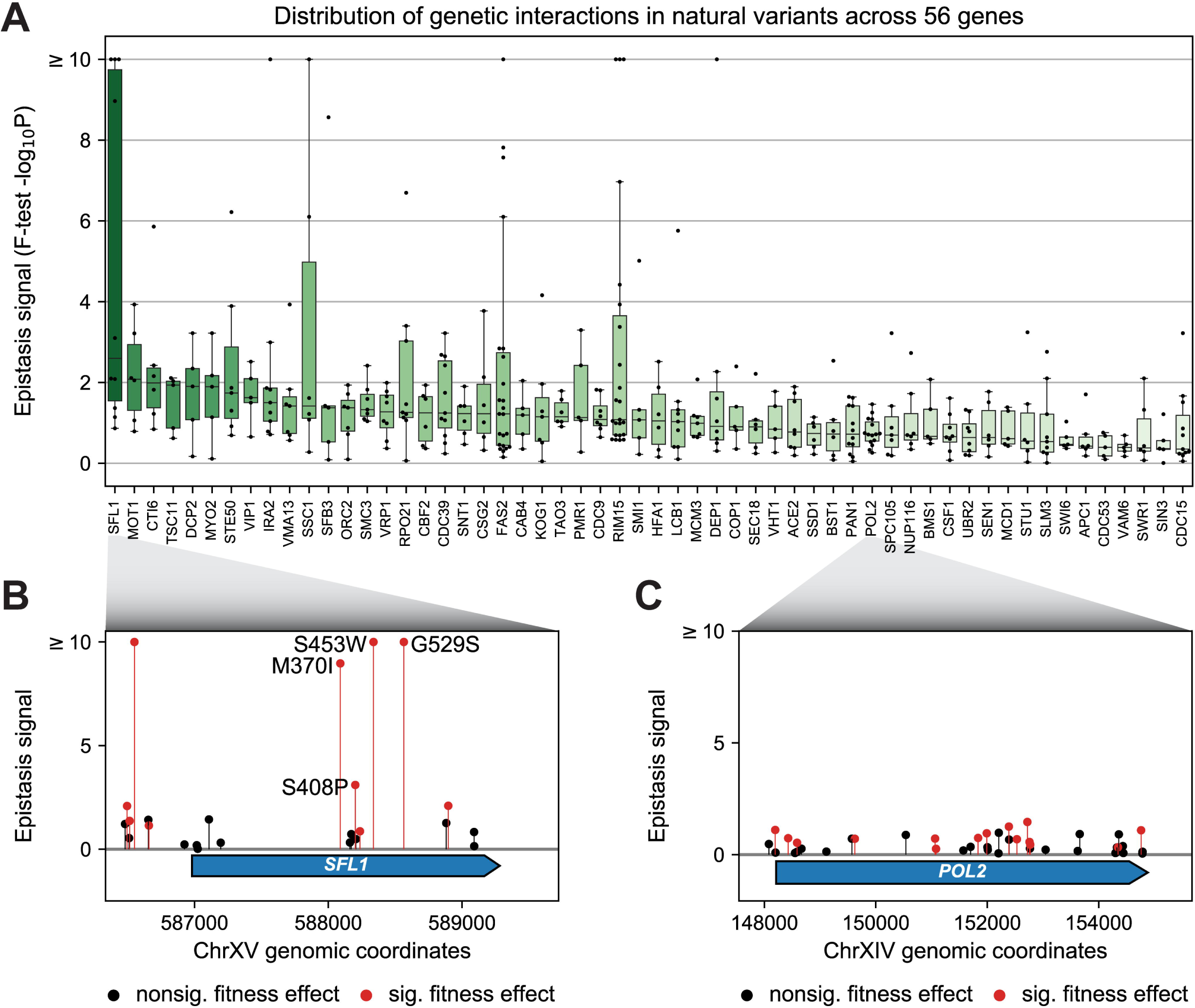
Genetic interactions are concentrated in key genes associated with differences in physiology between strains. (A) Genetic interactions are concentrated in a few key genes. Epistasis signal (F-test – log_10_[P-value]) reflects the extent to which genetic interactions modify the variant fitness effect. Variants with no significant effect in any strain are excluded. Color in boxplot reflects median –log_10_[P-value], from high (dark green) to low (light green). Large –log_10_P values are capped at 10 for visualization. (B) Multiple *SFL1* variants exhibit strong epistasis signal. Variants with significant fitness effects (red) are affected by genetic background more than variants with non-significant fitness effects (black). Amino acid changes of missense mutations with significant epistasis are marked. (C) In contrast, little epistasis signal is observed among *POL2* variants, regardless of the presence of significant fitness effects.

*SFL1* is a transcriptional repressor of genes involved in flocculation (Atsushi et al., 1989; Robertson and Fink, 1998) – the process where yeast cells form multicellular aggregates called flocs in response to changes in the microenvironment (Soares, 2011). Flocculation ability is a highly variable complex trait and is the most significant functional enrichment among open reading frames (ORF) missing from a subset of the 1,011 yeast strains collection (Peter et al., 2018). In our study, YPS and YJM are flocculent, while BY and RM are non-flocculent due to a *FLO8* mutation in BY (Liu et al., 1996) and a small deletion in *FLO1* in RM (Brem et al., 2002). Consequently, knocking out *SFL1* in the flocculent strains produces rapid flocculation and sedimentation upon exposure to calcium, but not in the non-flocculent strains (Figure 6A).

**Figure 6.**
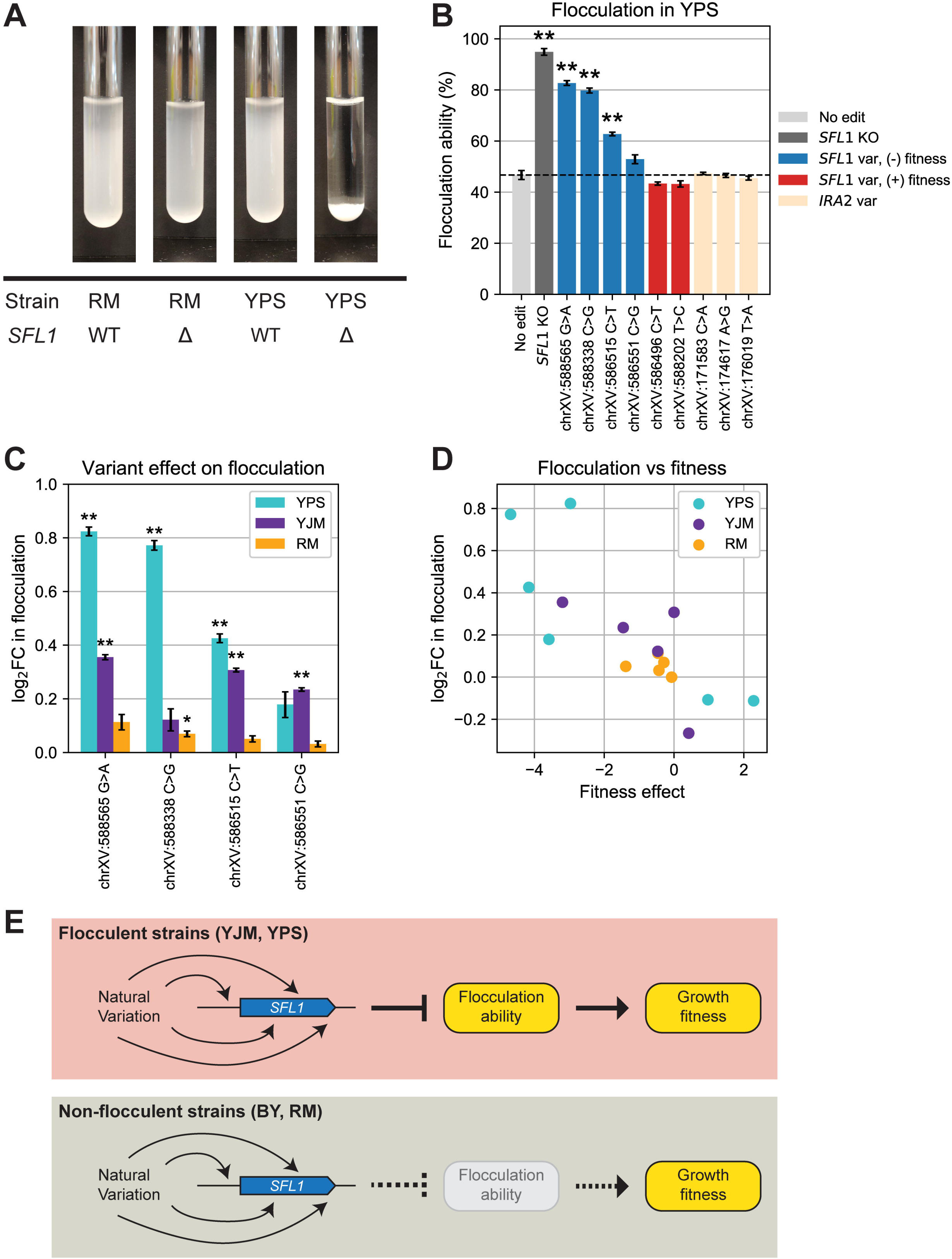
Fitness effects of *SFL1* natural variants are associated with strain-specific changes in flocculation ability. (A) Knocking out *SFL1* causes rapid flocculation in flocculent YPS upon exposure to calcium, but not in non-flocculent RM. WT: wild-type, Δ: knock-out mutant. (B) *SFL1* variants with negative fitness effects are associated with increased flocculation ability in YPS. (C) *SFL1* variants increase flocculation ability in a strain-specific manner. Data is represented by mean ± SEM. Asterisks indicate significance levels for t-test after Bonferroni adjustment. *p<0.05 **p<0.005 (D) *SFL1* variant fitness effects are negatively correlated with change in flocculation ability. (E) In flocculent strains, natural variation in *SFL1* modifies the regulation of flocculation ability and affects growth fitness directly and indirectly. However, non-flocculent strains weaken this relationship via loss of function in the intermediate phenotype.

To investigate whether *SFL1* variants that affect fitness also modify flocculation ability, we introduced the variants into each strain and measured the change in flocculation ability in each sequence-verified mutant relative to its parent strain. We found that *SFL1* variants that produce a negative fitness effect in YPS also increased flocculation ability, and vice versa (Figure 6B). To rule out the possibility that the flocculation change is caused by non-specific changes to strain fitness, we also tested variants in a gene not directly related to flocculation (*IRA2*) that have significant fitness effects. We found no change in flocculation ability, supporting the notion that the relationship between fitness and flocculation ability is *SFL1*-specific. In addition, the change in flocculation ability varies depending on genetic background; variants that increase flocculation ability in YPS and YJM have little to no impact in RM (Figure 6C). Furthermore, the degree of change in flocculation ability caused by the variant is negatively correlated with its growth fitness effect (Pearson’s r = -0.79, p < 10^-3^; Figure 6D), as expected if flocculation is mediating these fitness effects. In sum, our results suggest that *SFL1* variants may affect fitness via their effects on flocculation, thus providing a mechanistic explanation for why these fitness effects are greater in flocculent strains.

## Discussion

Our work reveals the importance of studying genetic interactions involving individual genetic variants to reveal molecular mechanisms governing complex traits. This was achieved through the high-throughput precision editing and lineage tracking capabilities of CRISPEY-BAR, enabling us to connect the genetic interactions of natural variants to complex traits and gain novel insight into their molecular underpinnings, which has not been possible in QTL studies or knockout screens. We presented evidence for how single nucleotide variants in *SFL1* mediate changes in growth fitness and flocculation ability in a genetic context-dependent manner. Similar strategies can be adopted to investigate genes implicated in other highly variable complex phenotypes. For instance, *RIM15* is a glucose-repressible protein kinase involved in cell proliferation in response to nutrient availability (Roosen et al., 2004; Su and Mitchell, 1993; Vidan and Mitchell, 1997) which we found to contain several variants with strong genetic interactions (Figure 5A). Given the diversity in nutrient types and quantity across ecological niches, we expect that *RIM15* will be differentially adapted in strains isolated from different environments and warrants further study.

The strong association between flocculation ability and growth fitness suggests that these traits are linked. We propose a mechanism to connect genotype to phenotype in *SFL1*: variants that reduce *SFL1*’s ability to repress flocculation genes lead to increased flocculation, which could reduce their growth fitness in the pool. However, strains with non-functional flocculation pathways disrupt this relationship, weakening the connection between *SFL1* variation and growth fitness (Figure 6E). This effect could reflect reduced growth rate within flocs, or perhaps that flocs are less likely to be included in serial dilutions (though no visible flocs were observed during the experiment). This does not rule out the possibility of *SFL1* affecting fitness through flocculation-independent pathways, particularly since *SFL1* interacts with and regulates other genes such as SUC2 and HSP30 (Ansanay Galeote et al., 2007; Song and Carlson, 1998) which may contribute to changes in growth fitness.

The variants screened in this study were selected to be in genes whose deletions exhibit a high number of pairwise genetic interactions (Costanzo et al., 2016), raising the question of how this ascertainment bias may have affected the trends we observed. We did not detect any significant association between a gene’s number of pairwise genetic interactions and the likelihood of its natural variants experiencing background effects, suggesting that these “hub genes” may not be enriched for epistasis of individual natural variants. Furthermore, the striking enrichment of epistasis among beneficial variants is unlikely to be a result of our variant selection strategy and therefore may be a general feature of epistasis in natural genetic variation.

This asymmetry in the distribution of genetic interactions across the range of fitness effects, where beneficial variants are more likely to exhibit strain-specific effects, leads us to hypothesize that strong strain-specific effects prevent these variants from being beneficial in all genetic backgrounds, leading to their maintenance as polymorphisms in the population. Another open question is whether our findings in yeast translate to other species. Although it is impractical to conduct genetic screens and high-throughput phenotyping in certain species such as humans, there are many tools available (Erwood et al., 2022; Hanna et al., 2021) to screen human cell lines derived from diverse populations to study complex traits at the cellular, tissue and organoid level.

Perhaps the most intriguing question raised by our results is why are beneficial variants so highly enriched for epistasis? Recently, the idea of “idiosyncratic epistasis”, where phenotypic effects depend heavily on the specific genetic background, has been proposed to explain trends such as diminishing returns of beneficial variants and increasing costs of deleterious variants (Bakerlee et al., 2022; Lyons et al., 2020; Reddy and Desai, 2021). This model provides a useful framework to explain our results. In this model, for a variant to be beneficial the sum of its interaction terms with all other genomic loci must be positive. However, in any well-adapted species the genetic interactions of most mutations will likely tend to be negative, resulting in a bias towards deleterious effects overall. In this framework, beneficial variants can be seen as outliers that have “won the epistatic lottery” and overcome the negative bias, but in these cases one would not expect the same variant to win this lottery in four diverse backgrounds. Therefore, beneficial variants would be more likely to have a higher variance in their fitness effects for the same reason that a lottery winner is expected to have higher variance in their payoff across multiple lottery tickets as compared to a lottery loser. Another prediction of idiosyncratic epistasis is that the DFE of each strain should be largely determined by its baseline fitness (Chou et al., 2011; Kryazhimskiy et al., 2014; Wei and Zhang, 2019); we did not observe this (Figure S1), though with only four strains we were not well-powered to observe this effect.

In conclusion, our study highlights the non-random distribution of genetic interactions: epistatic effects are enriched in beneficial variants and at intermediate allele frequencies. Despite only surveying a small fraction of all natural variation in yeast, we were able to accurately quantify the fitness effects of more than 1,800 natural variants and detect significant genetic interactions involving many of them. We expect that further studies of epistasis with single-nucleotide resolution will continue to reveal the extent to and molecular mechanisms by which epistasis shapes the genetic architecture of complex traits.

## Methods

### Construction of CRISPEY-BAR yeast strains

BY4742 (BY) was previously engineered to include Ec86 reverse transcriptase (RT) and SpCas9 via the integration of plasmid pZS157 (Sharon et al., 2018). To integrate RT and SpCas9 in non-BY yeast strains, pZS157 was modified via Gibson assembly to add a NatMX6 antibiotic resistance marker to the RT-SpCas9 integration cassette (pTEF::NatMX6::tTEF, pGAL10::Ec86RT::tGAL10, pGAL1::SpCas9::tCYC), flanked by 500 bp homology arms targeting *URA3* or *LYS2* to generate plasmids pRA027 and pRA028, respectively. pRA027 was double digested with restriction enzymes MfeI and SacII (New England Biolabs) to integrate RT-SpCas9 in YJM789 (YJM), and selected for nourseothricin resistance and uracil auxotrophy. pRA028 was double digested with restriction enzymes BstZ17I and SacII (New England Biolabs) to integrate RT-SpCas9 in RM11-1a (RM) and YPS128 (YPS), and selected for nourseothricin resistance and lysine auxotrophy. Functional validation of CRISPEY-BAR editing in the non-BY yeast strains was performed with plasmid pSAC212 (Chen et al., submitted) that inserts a nonsense mutation in *ADE2* – which turns yeast colonies pink in adenine-limiting conditions, as described by Sharon et al. (2018). Red colonies were counted from dilution plating during editing induction to determine editing efficiency, which was found to be similar across the four strains (Figure S9).

Next, a 6 bp strain-specific sequence was inserted 75 bp away from the CRISPEY-BAR barcode integration site in gene YBR209W via galactose induction of CRISPEY editing (Sharon et al., 2018) to facilitate identification of each yeast strain and avoid cross-contamination (Table S5). In addition, we switched the mating types of RM and YPS from MAT**a** to MATα using a modified version of the CRISPR-based protocol by Xie et al. (2018): RM and YPS yeast were grown in synthetic complete medium with 2% raffinose overnight before being grown to log phase in synthetic complete medium with 2% galactose for 4 hours to induce expression of RT and Cas9 prior to lithium acetate/PEG transformation (Gietz and Schiestl, 2007) with plasmids pXZX501 and NsiI-digested plasmid pXZX353, kindly provided by Zexiong Xie. Transformants were selected from plating on synthetic medium with uracil drop-out and 2% galactose, and successful mating type switch was verified by test crosses to MAT**a** strains BY4741 and growth in synthetic drop-out medium lacking lysine and methionine; only RM/YPS transformants that have switched to MATα could mate with BY4741 to form diploids that can grow without supplemental lysine and methionine.

### Media

CRISPEY-BAR pooled editing was performed with synthetic medium with uracil drop-out and 2% raffinose (SD-URA 2% raffinose), as well as synthetic medium with uracil drop-out and 2% galactose (SD -URA 2% galactose), previously described by Sharon et al. (2018). Growth competitions were performed in synthetic complete medium and 2% glucose (SCM: 1.7 g/L yeast nitrogen base without amino acids and ammonium sulfate (YNB), 5 g/L ammonium sulfate, 2 g/L Drop-out Mix Complete w/o Yeast Nitrogen Base, 2% glucose). Removal of plasmids post-editing was performed in synthetic medium with 5-fluoroorotic acid (SCM +5-FOA: 1.7 g/L YNB, 5 g/L ammonium sulfate, 2 g/L Drop-out Mix Complete w/o Yeast Nitrogen Base, 1 g/L 5-Fluoroorotic Acid Monohydrate).

### Selection of yeast natural variants for library

Natural variants were selected from the ∼1.75 million mutations documented in 1,011 yeast genomes (Peter et al., 2018) using a custom Python script (select_crispey3_library_variants.ipynb). 800 genes with a large number of genetic interactions based on previous work by Costanzo et al. (2016) were identified and natural variants in and around these genes (up to 500 bp away from CDS) were screened for their eligibility to design CRISPEY-BAR oligomers for editing: First, the four yeast strains must possess the reference allele at the variant locus to allow editing to the alternate allele. Second, variants had to be targetable by at least two guide RNA sequences within 6 bp of the variant so that multiple CRISPEY oligomers could be designed to assess fitness effects from independent guides. The variant’s proximity to the guide sequence ensures a high rate of incorporation via homology-directed repair (HDR) near the Cas9 cut site (Guo et al., 2018). To avoid overwriting existing polymorphisms in each yeast strain during CRISPEY-BAR editing, natural variants within 30 bp of existing RM, YJM and YPS polymorphisms were excluded. Several genes that may affect CRISPEY editing were also excluded from the variant search (see script for full list). Variants that are singletons or doubletons across the 1,011 yeast strains were also excluded. For polymorphic loci with multiple alternate alleles, the non-reference allele with the highest allele frequency was selected for editing. Of the 18,255 natural variants that passed the filtering criteria, 1,000 variants with allele frequencies greater than or equal to 2% – as defined in the 1,011 yeast genomes VCF file – spanning 103 genes were selected for the CRISPEY-BAR library in this work. In addition, a matching number of “very rare” variants (defined as variants with an allele count of 3 or 4 in the 1,011 yeast genomes VCF file) within each gene were also selected, bringing the total number of variants to 2,000. Selected variants for this work have the variant ID prefix “GXG” in the VCF file used for CRISPEY-BAR oligomer design.

### CRISPEY-BAR barcode-donor-guide oligomer design

Variants were provided to a custom Python script for designing CRISPEY-BAR oligomers (crispey3_design_library.ipynb), which is adapted from previous code by Sharon et al. (2018). The pipeline extracts all available guide sequences and assigns quality scores using Azimuth (Doench et al., 2016), as well as searches for off-targets for each guide using Bowtie2 (Langmead and Salzberg, 2012) with the same parameters as previously described. The off-target search was done in the genome assemblies of the four yeast strains in this study; the reference genome assembly used was version R64-1-1 (SacCer3) from the UCSC Genome Browser (https://hgdownload.soe.ucsc.edu/downloads.html#yeast). The RM (RM11-1A_SGD_2015_JRIP00000000.fsa) and YJM (YJM789_Stanford_2007_AAFW02000000.fsa) assemblies were obtained from the Saccharomyces Genome Database (http://sgd-archive.yeastgenome.org/sequence/). The YPS assembly was obtained from Yue et al. (2017) (http://yjx1217.github.io/Yeast_PacBio_2016/data/Nuclear_Genome/YPS128.genome.fa.gz). Each guide had a 108 bp asymmetric, strand-specific donor sequence designed with a 40:68 split in upstream and downstream bases around the Cas9 cut site; this facilitates the creation of multi-copy single-stranded DNA (msDNA) templates that have long 5’ and short 3’ homology arms, which have been demonstrated to improve HDR efficiency (Richardson et al., 2016). Additional small shifts in donor position were permitted to avoid excluded sequences such as 10 bp homopolymers, as well as NotI, SphI and AscI recognition sequences, while maintaining a minimum upstream length of 30 bp and minimum downstream length of 55 bp. In addition to the natural variants, 42 short deletion mutations that disrupt genes with known effects on growth in BY (Breslow et al., 2008) were included as controls (referred to as “ladder set” in this study) to assess the reproducibility of CRISPEY-BAR fitness measurements. The guides and donors used for the ladder set were obtained from Bao et al. (2018). Finally, we designed 74 non-editing guide-donor control sequences that represent wild-type fitness in growth competitions: 30 were previously used in Sharon et al. (2018), 42 were adapted from the ladder set by scrambling its guide and donor sequences, and two have guides targeting GFP sequence which is not present in these yeast genomes.

Subsequently, 12 bp programmed barcodes were assigned to each guide-donor combination to allow tracking of the variant edit by barcode sequencing in the growth competitions. The list of barcodes was generated by Chen et al. (submitted) using DNA-based Hamming codes (Bystrykh, 2012). For the guide-donor combinations corresponding to the 42 ladder set mutations and 74 non-editing controls, multiple programmed barcodes were assigned to provide additional replication in the library. The assigned barcode, donor and guide sequences were assembled into 250 bp CRISPEY-BAR oligomers (design described in Chen et al. (submitted)) and sorted into groups of 121 for matrixed oligo pool synthesis by Twist Biosciences. In total, 5,319 oligomers (4,957 targeting natural variants, 181 non-editing controls, 181 targeting ladder set mutations) were synthesized for use in this study (Table S1).

### Yeast CRISPEY-BAR library cloning

Oligomers were cloned into CRISPEY-BAR vectors as described in Chen et al. (submitted). Oligomers in each well of a 384-well plate were amplified with Q5 hot-start DNA polymerase (New England Biolabs) using primers SAC615 and SAC576, and pooled by equal DNA concentration. The pooled oligos were next assembled into the NotI-HF-digested plasmid pSAC200 using NEBuilder HiFi DNA Assembly Cloning Kit (New England Biolabs) and transformed via electroporation into Endura Electrocompetent *E. coli* cells (Lucigen) using a MicroPulser Electroporator (Bio-Rad). Transformed bacteria were plated on LB with Carbenicillin plates and incubated at 32°C for >20 hours, and the bacterial lawn later scraped for plasmid extraction using NucleoBond Xtra Midi Plus (Takara Bio). The plasmid pool was then triple-digested with SphI-HF, AscI, NotI-HF and treated with CIP (New England Biolabs), followed by magnetic bead purification with NucleoMag NGS Clean-up and Size Select (Takara Bio). To prepare the CRISPEY-BAR middle insert for ligation, the ∼400 bp fragment was amplified by Q5 PCR with one reverse primer (SAC590) and one of six distinct forward primers (SAC591, 592, 594, 596, 603, 604), using the plasmid pSAC212 as a template. The ligation insert appends one of six possible 9 bp unique molecular tags to each programmed barcode, which provides additional technical replication to each barcode in the library. The inserts were pooled at equal concentration and double-digested with SphI-HF and AscI, cleaned up with NucleoMag NGS Clean-up and Size Select beads (Takara Bio), and ligated with the aforementioned triple-digested plasmid pool using T4 DNA ligase (New England Biolabs) at 16°C overnight with a insert:vector molarity ratio of 20:1. The ligation reactions were heat-inactivated at 65°C for 10 minutes and cleaned up with NucleoMag NGS Clean-up and Size Select beads (Takara Bio) before electroporation into Endura Electrocompetent *E. coli* cells (Lucigen) using a MicroPulser Electroporator (Bio-Rad). Transformed bacteria were plated on LB with Carbenicillin plates and incubated at 32°C for >20 hours, and the bacterial lawn later scraped for plasmid extraction using NucleoBond Xtra Midi Plus (Takara Bio). 20 individual bacterial colonies were separately picked from a dilution plating and miniprepped using Nucleospin Plasmid (Takara Bio), and the extracted plasmids were Sanger sequenced to verify the cloning quality of the library. 19 clones were found to have the full-length oligomer with the correct middle insert ligation necessary for functional editing and barcoding with CRISPEY-BAR, while 1 had a short insert ligation which is predicted to be non-functional in the pool. Only plasmids that have been successfully assembled are able to conduct barcoding of cells during CRISPEY-BAR pooled editing; even if present in a pooled competition, cells without barcode edits will not be sequenced or affect the quality of data collected.

The post-ligation plasmid pool was transformed into each yeast strain separately, using the lithium acetate/PEG method (Gietz and Schiestl, 2007). Yeast grown overnight in liquid YPD at 30°C were transferred to fresh YPD with starting OD_600_ of ∼0.3 and grown at 30°C to log phase over 4-5 hours. 12-25 ug of plasmid was transformed into each yeast strain, with the aim of getting ∼10^6^ transformants in each yeast library. This was achieved by heat shock at 42°C for 25 minutes for BY, RM and YJM, and 40°C for 40 minutes for YPS. In addition, another replicate transformation of BY was performed with the same plasmid pool to generate two BY libraries that were independently edited and competed to assess the reproducibility of fitness measurements across independent experiments. Yeast transformants were selected for on synthetic media with uracil dropout (SD -URA) 2% agar plates and incubated at 30°C for 48 hours. Transformation efficiency was estimated by dilution plating of the transformed yeast: 1-15 million transformants were recovered for each of the 5 yeast libraries prepared (two for BY, one for RM, YJM and YPS). Yeast transformant lawns were scraped off plates after 48 hours and frozen down in 15% glycerol stocks at -80°C for future CRISPEY-BAR editing.

### CRISPEY-BAR pooled editing

CRISPEY-BAR pooled editing was done similarly to previous work by Sharon et al. (2018): Each yeast library was thawed from frozen glycerol stock and added to 150 mL SD -URA 2% raffinose, with approximately 10^9^ cells, and incubated at 30°C shaking at 250 rpm for 24 hours. To begin galactose induction of editing, 5x10^8^ cells of pre-edited yeast were added to 150 mL SD -URA 2% galactose, with a starting OD_600_ = 0.3, and incubated at 30°C shaking at 250 rpm. Editing lasted for 72 hours, with the yeast libraries being transferred to 150 mL of fresh SD -URA 2% galactose every 24 hours. 5x10^8^ cells were moved at every transfer, with a mean library coverage of ∼91,000 cells per guide-donor. All libraries were edited for ∼16 generations, apart from YJM which only grew 9 generations due to a defect in *GAL2* which prevents YJM from metabolizing galactose anaerobically (Wei et al., 2007). Despite this, all strains showed similar levels of editing after 72 hours of induction (Figure S9). To remove CRISPEY-BAR plasmids after editing, 5x10^8^ cells from each edited yeast library were added to 150 mL SCM and incubated at 30°C shaking at 250 rpm for 24 hours to allow relaxation of plasmid selection. After ∼5.5 generations of non-selective growth, 5x10^8^ cells were transferred to 150 mL SCM +5-FOA and incubated at 30°C shaking at 250 rpm for 24 hours to select against the *URA3*-bearing CRISPEY-BAR plasmid over ∼5.5 generations. Dilution plating on SD -URA plates was used to determine the number of URA+ cells left after 5-FOA selection; the mean proportion of plasmid-bearing cells remaining in the libraries is ∼0.1%. The edited, plasmid-free yeast libraries were frozen down in 15% glycerol stocks at -80°C in preparation for growth competitions.

### Growth competition

Each yeast library was thawed from frozen glycerol stock and added to 150 mL SCM, with approximately 10^9^ cells, and incubated at 30°C shaking at 250 rpm overnight. Three biological replicate growth competitions were seeded from the same starting culture in separate flasks containing 150 mL fresh SCM, with an initial OD_600_ = ∼0.13 (approximately 2x10^8^ cells). Growth competitions were incubated at 30°C shaking at 250 rpm for 8 hours, and the first time point (0^th^ generation) was recorded at the end of 8 hours. During this and every subsequent 8-hourly time point, 2x10^8^ cells from each growth competition flask were transferred to 150 mL fresh SCM to return the competition culture to OD_600_ = ∼0.13, and then incubated at 30°C shaking at 250 rpm for 8 hours to let competition cultures divide for ∼5 generations, reaching a final OD of ∼4-6. The growth competitions lasted 96 hours, for a total of ∼60 generations (Table S6). We collected samples from the competition cultures at every time point by centrifuging ∼5x10^8^ cells and removing the supernatant, storing the yeast pellets at -20°C for later genomic DNA extraction and barcode sequencing.

### Barcode extraction and sequencing

Yeast pellets from the 0^th^, 24^th^, 48^th^, 72^nd^ and 96^th^ hour of growth competitions (corresponding to approximately 0^th^, 15^th^, 30^th^, 45^th^ and 60^th^ generations, respectively) had their total genomic DNA extracted using MasterPure Yeast DNA Purification Kit (Lucigen), with cell lysis solutions and RNAse A treatment volumes scaled up 4x to adequately lyse all cells and remove RNA, based on manufacturer’s recommendations. Genomic DNA extract concentrations were quantified using Qubit dsDNA HS kit (ThermoFisher). CRISPEY-BAR barcodes integrated in the genome were amplified with a first step PCR in 8 tubes of 50 uL reactions using Q5 hot-start DNA polymerase (New England Biolabs) following manufacturer recommendations, each containing 1.5 ug genomic DNA, 1 uM of forward primer (SAC261), 1 uM of a mix of 8 staggered reverse primers (SAC327-334) and supplemented with 1 M betaine. PCR thermocycler settings were as follows: initial denaturation of 98°C for 2 min, followed by 18 cycles of 98°C for 10 s and 65°C for 20 s, and a final extension at 72°C for 2 min. The staggered primers provide additional sequence complexity for read 2 during barcode sequencing. In addition, the primers used in this PCR carry partial Illumina adaptor sequences that permit indexing with Illumina dual-indexing primers in the second step PCR: after cleaning up with NucleoSpin Gel and PCR Clean-Up (Takara Bio), amplified barcodes were indexed using Q5 hot-start DNA polymerase (New England Biolabs) following manufacturer recommendations, with 1 uM i5 index primer (S502-503, S505-508, S510-511, S513, S515-518, S520-521) and 1 uM i7 index primer (N701-707, N710-712, N714-716, N718-724). PCR thermocycler settings were as follows: initial denaturation of 98°C for 2 min, followed by 7 cycles of 98°C for 10 s and 70°C for 20 s, and a final extension at 72°C for 2 min. The indexed barcodes were cleaned up with NucleoSpin Gel and PCR Clean-Up (Takara Bio) and DNA concentration measured using Qubit dsDNA HS kit (ThermoFisher). Samples were pooled by equal DNA concentration and size-selected with E-Gel SizeSelect II 2% agarose gel (Invitrogen), and elute was concentrated with NucleoSpin Gel and PCR Clean-Up (Takara Bio). The pooled samples were sequenced using 5 lanes of Illumina Nextseq 550 High-Output, using a custom read 1 primer (SAC354) and custom dual index sequencing (12 read 1 cycles, 64 read 2 cycles), with each growth competition triplicate sequenced to an average depth of ∼400 million reads. All primer sequences are provided by Chen et al. (submitted) and summarized in Table S7.

### Variant fitness effect estimation

Sequencing reads were processed and counted to assemble the barcode count matrix, as described in the script crispey3-count-barcodes.ipynb. After inspecting fastq files with FastQC (https://www.bioinformatics.babraham.ac.uk/projects/fastqc/), read 2 was trimmed using Cutadapt (Martin, 2011) with parameters “-g GGCCAGTTTAAACTT…GCATGGC -j 20 -e 0.2 --nextseq-trim 20 --discard-untrimmed -m 12 --pair-filter=first” to extract the 27 bp barcode sequence, consisting of 12 bp of programmed barcode from the library design and 9 bp of unique molecular tag introduced in the ligation step of library cloning, joined by a 6 bp linker. Read 1 was merged with trimmed read 2 using FLASH (Magoc and Salzberg, 2011) with parameters “-m 8 -M 12 -x 0.25 -t 20” to use the higher quality base for the programmed barcode sequence. Merged reads with no ambiguous bases were counted to assemble a barcode counts matrix, and each sequence was mapped to a barcode ID using an error-tolerant search (up to 1 mismatch) of all barcodes in the library design table to identify the associated guide and variant in the competition (Table S8)

Each barcode’s log fold change in abundance per generation was estimated with DESeq2 (Love et al., 2014), documented in the script crispey3-deseq2.ipynb. The number of generations (or culture doublings) elapsed in the growth competitions was estimated from OD_600_ measurements taken at every time point, which was fitted into the following linear model based on Sharon et al. (2018):

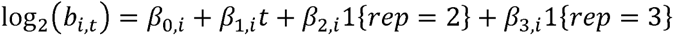

Where b_i,t_ is the normalized count of barcode *i* at generation *t*, and *t* is the number of generations from the start of the growth competition. The model accounts for different initial barcode abundances in each competition replicate flask (β_2_, β_3_), and calculates the average log fold change per generation (β_1_) of a barcode’s relative abundance across the competition triplicates. First, barcodes with fewer than 1000 counts across all 15 samples in a growth competition (5 time points, 3 replicates) were filtered out, and oligomers represented by fewer than 3 of the 6 possible unique molecular tags were excluded from downstream analyses. We first run DESeq2 with default settings on the neutral barcodes set, defined as the barcodes associated with the 181 non-editing oligomers in the library, to inspect the distribution of log fold change values and remove outliers with disproportionate leverage on the log2(fold-change)-baseMean relationship, where baseMean represents the average barcode count across all competition samples. DESeq2 was then run with default settings on all barcodes to inspect the library-wide distribution of log fold change values; we observed a small number of barcodes (<0.5% of the library) that had large positive log fold changes, defined as >3.5 standard deviations greater than the library-wide mean log fold change. Since these barcodes were not associated with specific guides/variants (i.e. the positive effect was not shared with other barcodes mapping to the same oligomer), we excluded these outliers from the data. Finally, the filtered barcodes were run with DESeq2 using the neutral barcodes set to estimate size factors; the barcode log fold changes now represent change relative to the neutral fitness distribution in the competition.

Individual barcode log fold changes were combined to estimate the variant fitness effect through a weighted least squares model using a custom Python script (crispey3-measure-fitness.ipynb). First, for each oligomer, we applied a robust outlier detection on its associated barcodes to remove ones with large median absolute deviations (MAD) from the median log fold change or counts (logFC > 3.5xMAD from median, baseMean > 5xMAD from median), since we expect that the unique molecular tags ligated to the oligomer barcode during library cloning should have relatively similar coverage and effect in the library. Next, we standardized the log fold changes across different growth competitions by calculating a Z-score for all barcodes based on the mean and variance of the neutral barcodes’ log fold change distribution in each growth competition; we refer to this score as the “fitness effect” in this study. To account for heteroscedasticity in the fitness effects of barcodes with different counts, we used the neutral distribution to calculate inverse-variance weights for each oligomer based on its median barcode’s average count in the competition (baseMean). Finally, barcode fitness effects and weights from each growth competition were jointly fitted into a weighted least squares model to calculate the variant fitness effect in each yeast strain. The model takes the variant, strain, and their interaction terms as independent variables, summarized in the form:

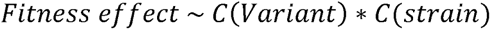

Where “C(variant)” and “C(strain)” are the variant and yeast strain categorical variables represented by dummy encoding in the model. Therefore, the variant fitness effect in a yeast strain is the difference between the weighted mean of fitness effects for barcodes associated with the variant and the weighted mean of fitness effects for neutral barcodes in that strain. The model also determined if the variant fitness effect is significantly different from neutral through a weighted t-test. The p-values were adjusted for multiple testing using the Benjamini-Hochberg procedure (Benjamini and Hochberg, 1995) and significant fitness effects were controlled at FDR = 0.1 (Table S2).

### Detecting significant genetic interactions in variants

The weighted least squares model allows tests for significant differences in a variant’s fitness effects between yeast strains through a weighted t-test; all pairwise differences in fitness effects between strains (e.g. six pairwise differences for variants measured in four strains) were calculated for variants with fitness measurements in at least two strains. The p-values were adjusted for multiple testing using the Benjamini-Hochberg procedure (Benjamini and Hochberg, 1995) and significant differences were controlled at FDR = 0.1 (Table S3). In addition, we tested a joint linear hypothesis using the F-test to find variants that have significant strain interaction terms that suggest significant differences in variant fitness effect in any of the strains. The F-test p-values were adjusted for multiple testing using the Benjamini-Hochberg procedure (Benjamini and Hochberg, 1995) and significant differences were controlled at FDR = 0.05 (Table S4). We consider a variant to be “epistatic” – more specifically, exhibiting significant genetic interactions due to genetic background effects – if it fulfilled both of the following criteria: 1. It passes a significant F-test with FDR < 0.05, and 2. It has at least two significant pairwise differences in fitness effects between strains with FDR < 0.1. If a variant’s fitness effect has only been estimated in two strains, we count it as epistatic if it has a significant F-test and pairwise fitness difference between the two strains available. We require two significant pairwise differences to reduce the number of false positives arising from variability in fitness effects between experiments. In practice, these criteria controlled the false positive rates at less than 5% when we tested non-editing oligomers in the library, which are expected to have no epistasis.

### Validation of pool effects

Twenty guide-donor oligomers from the library (19 targeting natural variants and one non-editing neutral) were synthesized as gene fragments by Twist Biosciences. 16 were synthesized as full length oligos, while four were amplified via Q5 PCR (New England Biolabs) with forward primer RA199 and oligo-specific primers to append the guide sequence (primers RA213-216) and the 3’ homology arm (primers RA209-212) to obtain each full-length oligomer. Each oligomer was assembled into the CRISPEY-BAR vector following the same procedure as the library cloning. Individual colonies had their plasmids extracted using NucleoSpin Plasmid (Takara Bio) and Sanger sequenced to verify successful ligation with no sequence errors. The sequence-verified plasmids were individually transformed into the four yeast strains, and individually edited via galactose induction following the same procedure as the library editing. Dilution plating was performed to select individual colonies to genotype for successful barcoding and variant editing, before plasmids were removed via 5-FOA selection, as described for the pooled library. The barcoded, sequence-verified mutants were frozen in 15% glycerol stocks at -80°C. Separately, GFP was integrated into the parental strains by transforming them with HindIII-digested plasmid pGS62, previously described by Kao and Sherlock (2008). A total of 80 sequence-verified mutants (of which 25 had significant fitness effects in the pooled competitions) and four GFP-fluorescent wild-type strains were generated to validate pool fitness effects via pairwise growth competitions between wild-type and mutant strains, using the change in ratio between fluorescent and non-fluorescent cells over time to estimate the relative growth fitness of mutants.

Single colonies from each mutant and its fluorescent parentals were grown in SCM at 30°C overnight, and their OD measured on a TECAN microplate reader. Each pairwise competition was setup by mixing a non-fluorescent mutant strain with its fluorescent parental strain at a 2:1 ratio in 400 uL SCM, with each competition performed in duplicates in 96 deep-well plates. GFP-only and non-GFP-only controls were included in each plate. The plates were incubated at 30°C with shaking to allow cells to divide, and diluted ∼1:30 at every 8-hourly time point by transferring 13.3 uL of yeast to 400 uL fresh SCM, similar to the pooled growth competition setup previously described. OD measurements were taken on a TECAN microplate reader at the start and end of each time point to estimate the number of co-culture doublings (generations) for each pairwise competition. At the 0^th^, 24^th^, 48^th^, 72^nd^ and 96^th^ hour time points, cells were diluted 1:10 in phosphate-buffered saline solution and had fluorescent/non-fluorescent cell counts measured with Attune NxT Flow Cytometer, using the same instrument settings as Pothoulakis and Ellis (2018). Up to 50,000 events were counted for each time point sample. Events with forward scatter (FSC) below 25 x 10^3^ A.U., or FSC reaching maximum value were excluded from the count, in addition to secondary gating on the side scatter (SSC) to remove other non-yeast events. Gating for counting GFP-fluorescent and non-fluorescent cells was determined using GFP-only and non-GFP-only control samples; both populations were fully separable on the 530 nm band-pass filter BL1-A (area) and FSC-A (area) channels

Relative growth rates of mutants in pairwise competitions were calculated based on methods in Breslow et al. (2008). For each pairwise competition, the ratio of non-fluorescent to fluorescent (mutant to wild-type) cells was calculated for every sample and used to calculate the number of wild-type doublings from the number of co-culture doublings estimated from OD measurements, and the numbers were fitted in a weighted least squares model with the formula:

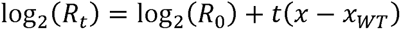

Where t is the number of wild-type generations, R_0_ and R_t_ are the mutant-to-wild-type ratios at the start and at time point t, respectively, and x and x_WT_ are relative growth rates of the mutant and wild-type strain, respectively. A replicate term was also included to account for different starting ratios between replicates, and weights are given by the square root of the number of events counted for each sample. In addition, samples depleted of mutant cells (ratio < 0.002) were excluded, resulting in one validation mutant being dropped due to having insufficient samples. Finally, the relative growth rates of mutants were normalized within each yeast strain such that the median growth rate is 1, to account for differences in the fitness of the GFP reference in each yeast strain. This is done with the assumption that the median validation mutant fitness is indistinguishable from neutral, which is a reasonable assumption since the median validation mutant of each strain has an approximately neutral fitness effect in the pooled competitions.

### Multiple logistic regression for factors associated with genetic interactions

All variant and gene properties were fitted into a multiple logistic regression to find factors with significant association with the incidence of epistasis due to genetic background, classified as present or absent in each variant based on the aforementioned criteria for detecting significant genetic interactions. The categorical variable for determining whether a variant is beneficial or deleterious is based on the sign of the median fitness effect across all four strains. When modeling allele frequency (AF), we used a categorical variable to separate common (AF ≥ 5%) and rare variants (AF < 5%), in addition to the AF continuous variable, to model the non-linear relationship between AF and epistasis incidence. Continuous variables were standardized prior to fitting the logistic model. Gene properties were ultimately excluded from the model as none significantly improved the model fit.

### Principal components analysis of population structure

Ecological origin information of the 1,011 yeast strains was obtained from Peter et al. (2018). For each natural variant in the CRISPEY-BAR library, we counted the number of strains carrying the variant and which ecological origins they represent, assembling a count matrix of the ecological origins represented for each variant. The count matrix was standardized, followed by linear dimensionality reduction via singular value decomposition into 15 principal components explaining 95% of variance. The top principal components separated the distribution of ecological origins among the natural variants (Figure S7), giving an approximate representation of the population structure for the variant library. The first six principal components explaining 74.8% of variance were added as covariates to the multiple logistic regression model to determine association with the incidence of genetic interactions in variants (Figure S8A).

### Flocculation assay

Flocculation measurements were taken from sequence-verified mutants based on methods by Bony et al. (1997): Mutants grown in YPD overnight were deflocculated by washing with 50 mM citrate (pH 3.0) 5 mM EDTA buffer twice and resuspended at ∼10^8^ cells/ml. Calcium chloride was added to the suspension to a final concentration of 20 mM to induce flocculation, and 3 mL of yeast was placed in 75 mm culture tubes. Tubes were incubated for 5 min at 30°C shaking at 100 rpm, and then left to stand vertically for 5 min. Samples were taken by pipetting 100 uL of yeast suspension from just under the meniscus before calcium induction and at the end of 5 min of standing post-induction, and OD_600_ measured by spectrophotometer. Flocculation ability was calculated by the formula:

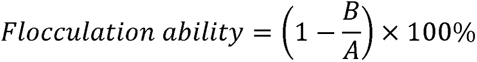

Where A is the OD measured of the fully deflocculated suspension and B is OD measured after 5 min of floc settling.

### Data and code availability

Raw sequencing data can be found at NCBI Sequence Read Archive, under BioProject accession number PRJNA826080. The code used for variant selection, oligomer design and processing of sequencing reads to estimate variant fitness effects are available on https://github.com/roy-ang/crispey3-epistasis. Scripts for generating figures are available by request.

## Acknowledgements

We thank M.M. Desai, D.A. Petrov and members of the Fraser lab for helpful comments on the manuscript; Z. Xie for providing plasmids to conduct the yeast mating type switch; V.K. Chen for providing GFP plasmid pGS62 and guidance on flow cytometry experiments; and S. Yeo for logistical support in the growth competitions. This work was supported by NIH R01 grants (GM097171, GM134228). R.M.L.A. was supported by the National Science Scholarship, from the Agency of Science, Technology and Research (A*STAR). S.A.C. was supported by Bio-X Stanford Interdisciplinary Graduate Fellowship, NIH grant 1F31ES030282 and NIH 5T32GM007276-42. A.F.K. was supported by the NIH NHGRI Stanford Genomic Training Program (5T32HG000044)

## Author contributions

H.B.F. conceived the project. R.M.L.A. designed the CRISPEY-BAR oligonucleotide library, did the editing and growth competition experiments, as well as data analysis for this project. S.A.C. developed the CRISPEY-BAR technology. Y.X. assisted with library cloning. A.F.K. generated the programmed barcodes and assisted in the development of fitness effects estimation scripts. R.M.L.A. and H.B.F. wrote the manuscript.

## Supplemental information

• Table S1. Guide-donor oligonucleotides designed in CRISPEY-BAR library.
• Table S2. Growth competition variant fitness effects in four yeast strains.
• Table S3. Pairwise differences in variant fitness effects between strains.
• Table S4. F-test results for differences in variant fitness effects across four yeast strains.
• Table S5. Yeast strains genotype information, including 6 bp strain-specific tag sequences.
• Table S6. Number of generations elapsed in each growth competition time point sample.
• Table S7. List of primers and validation oligonucleotides used in this project.
• Table S8. Barcode count matrix of all samples across growth competitions.

## Supplementary figure legends

**Figure S1.**
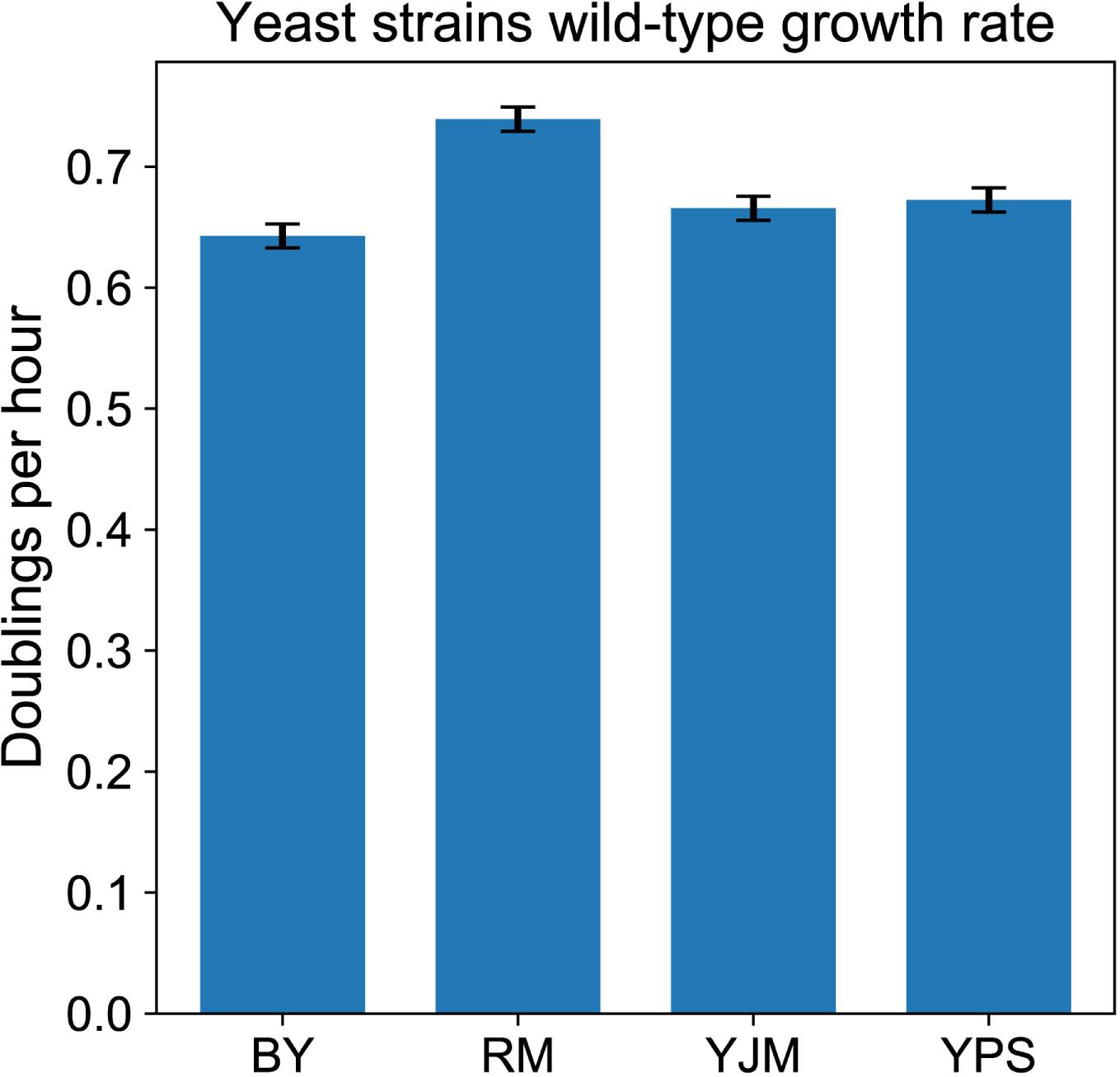
Wild-type growth rate of yeast strains in synthetic complete medium. Growth rate was calculated from number of doublings estimated from OD_600_ measurements over an 8-hour growth cycle, based on the growth competition setup and performed in triplicate per strain. Data is represented by mean ± SEM.

**Figure S2.**
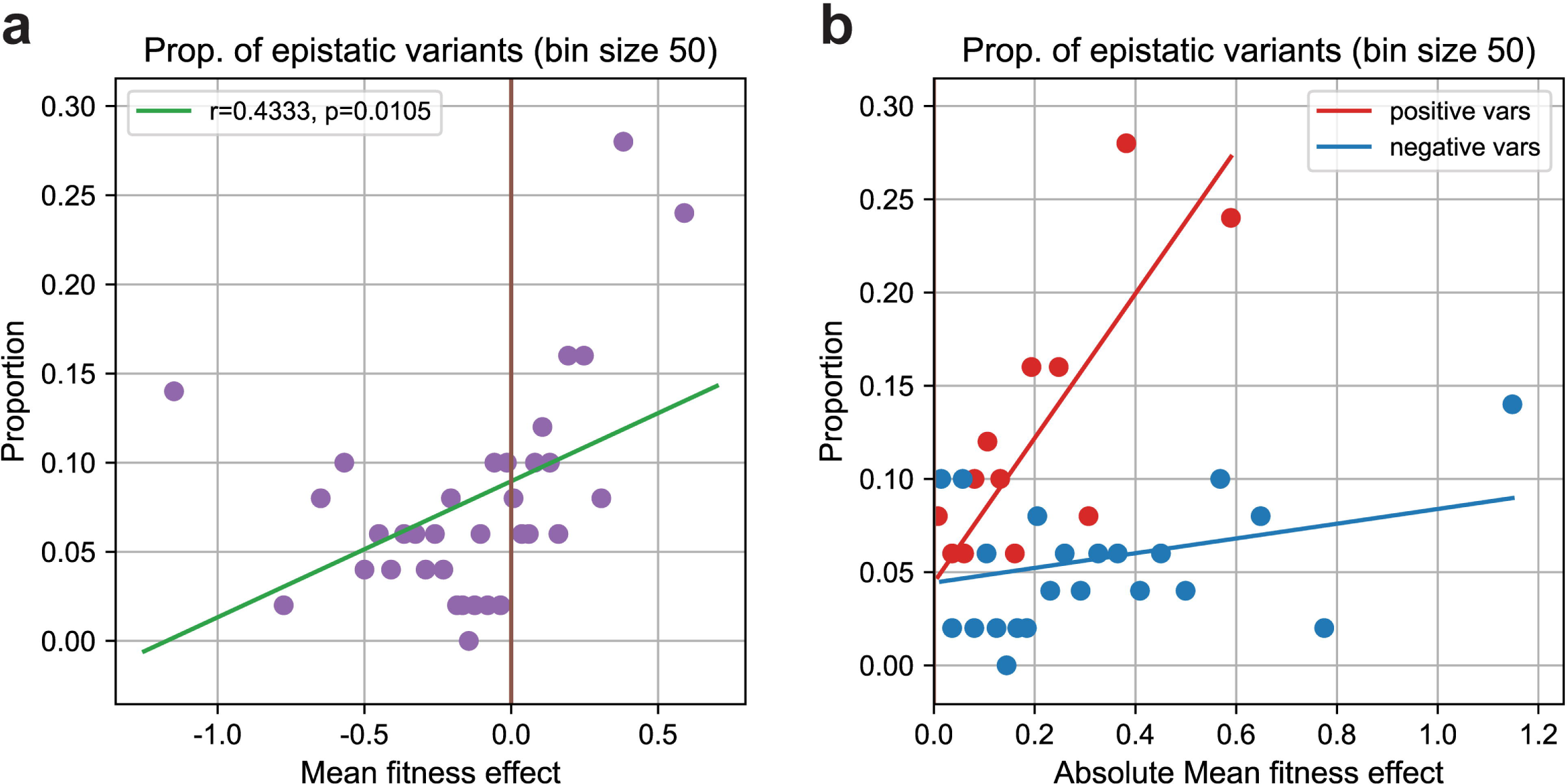
Proportion of epistatic variants increases with greater mean fitness effects. (a) Proportion of variants with genetic interactions increases with greater mean fitness effect. Each point represents 50 variants binned by increasing four-strain mean fitness effect. Green line represents the least squares fit. (b) Proportion of variants with genetic interactions is greater among variants with positive mean fitness effect than those with negative mean fitness effect, after controlling for effect size. Red and blue lines represent least squares fit for variants with positive mean fitness effect and negative mean fitness effect, respectively. Slopes are significantly different (least squares interaction coefficient p < 10^-7^).

**Figure S3.**
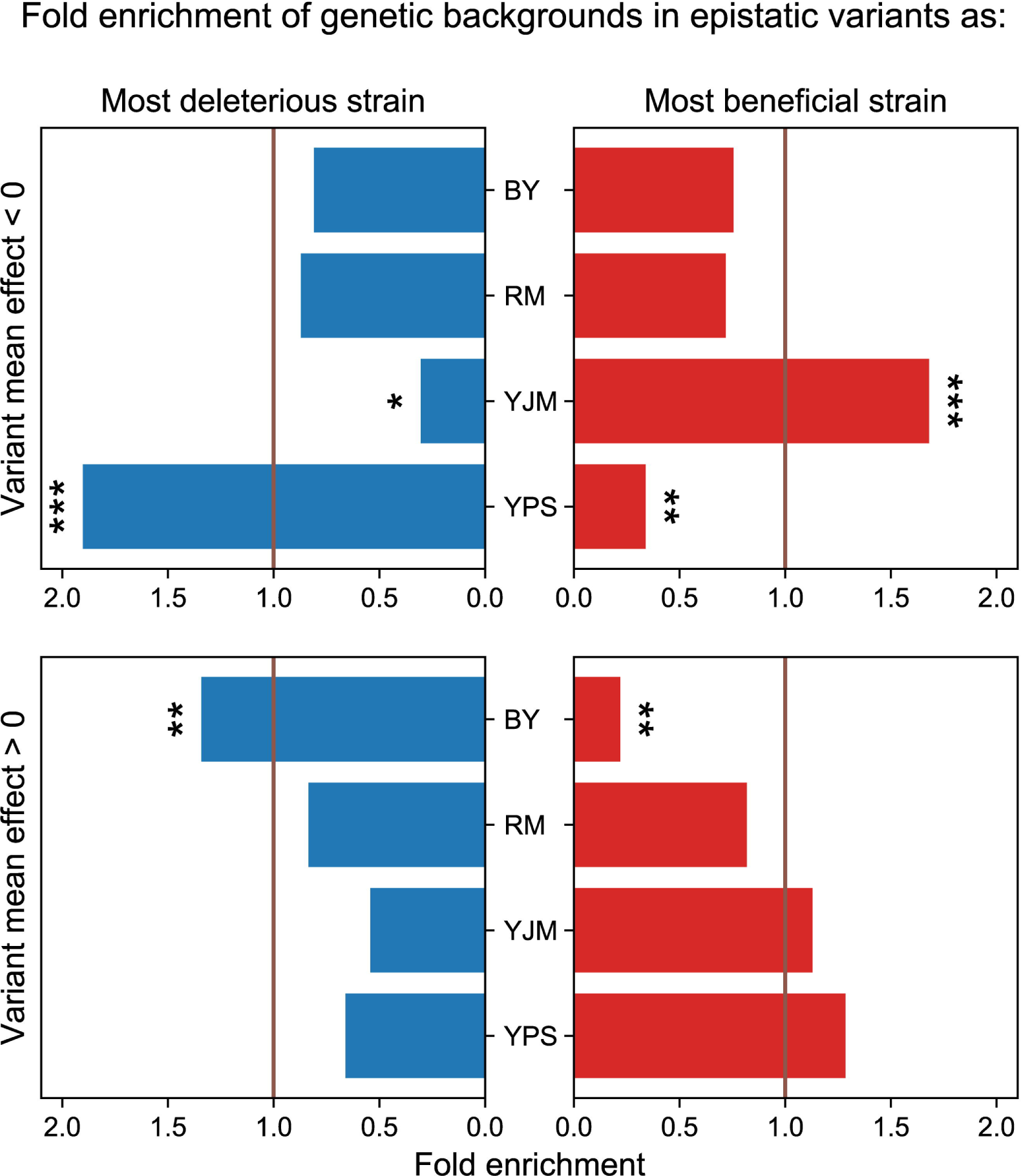
Genetic backgrounds are differentially enriched for having the most beneficial or deleterious effect among epistatic variants. Fold enrichments of each genetic background as having the most deleterious (left) or beneficial (right) effect in epistatic variants, relative to the library-wide average. Strains are differentially enriched between epistatic variants with negative mean fitness effect (top) and those with positive mean fitness effect (bottom). Asterisks indicate significance levels for Fisher’s exact test. *p<0.05 **p<0.005 ***p<0.0005

**Figure S4.**
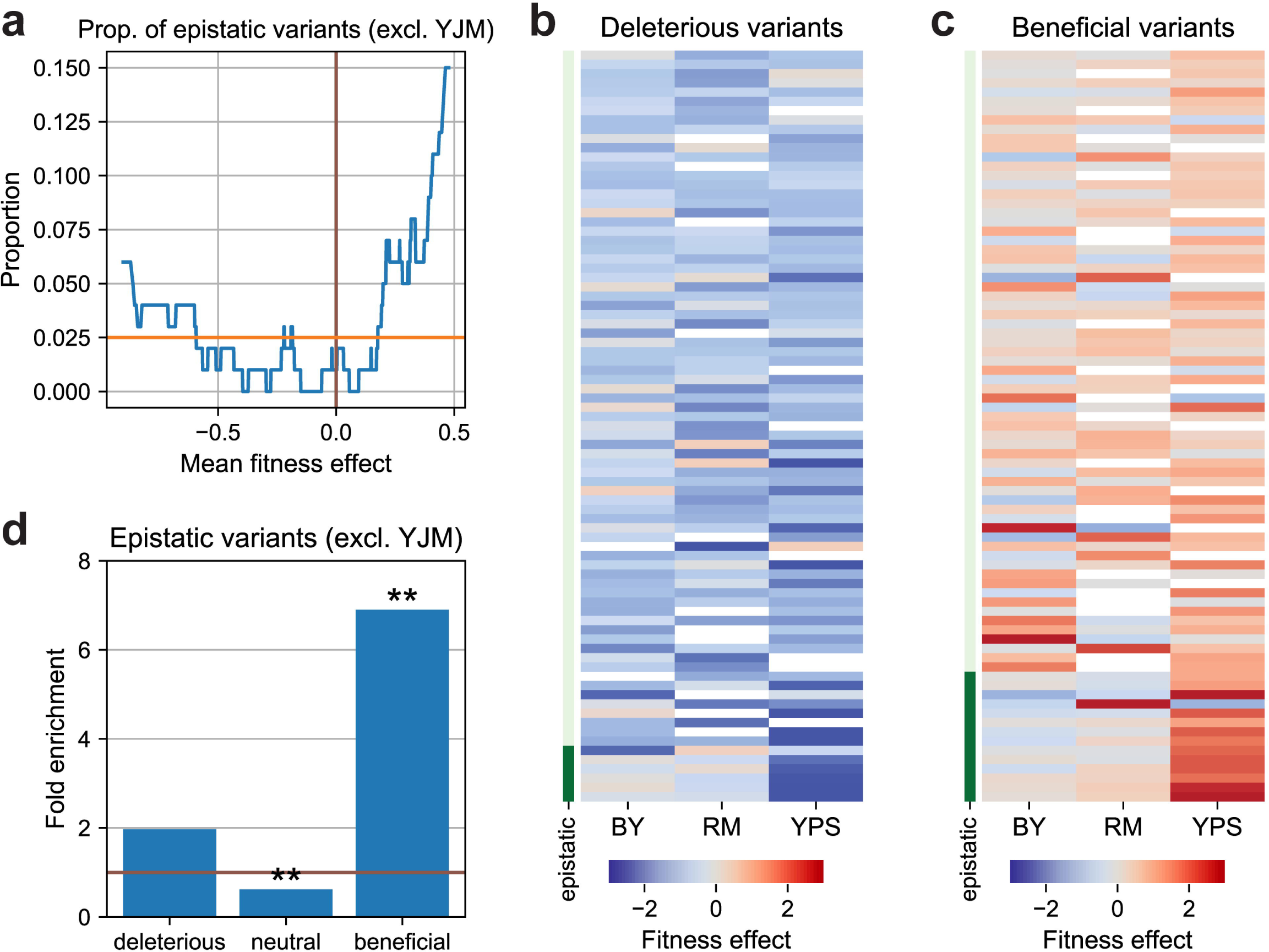
Enrichment of genetic interactions among beneficial variants without YJM. All panels are a reproduction of Figure 3A-D, but with YJM data excluded when calling genetic interactions in variants. Despite identifying fewer epistatic variants due to lower statistical power, the beneficial variants enrichment for epistasis remains significant. (a) Proportion of variants with genetic interactions increases with greater mean fitness effect. (b) Deleterious variants show consistent negative effects across yeast strains. (c) Beneficial variants are more likely to exhibit strain-specific fitness effects. Row colors on Y-axis indicate the presence (dark green) or absence (light green) of genetic interactions. Missing values are labeled white. (d) Genetic interactions are strongly enriched in beneficial variants. Asterisks indicate significance levels for Fisher’s exact test. *p<0.05 **p<0.005

**Figure S5.**
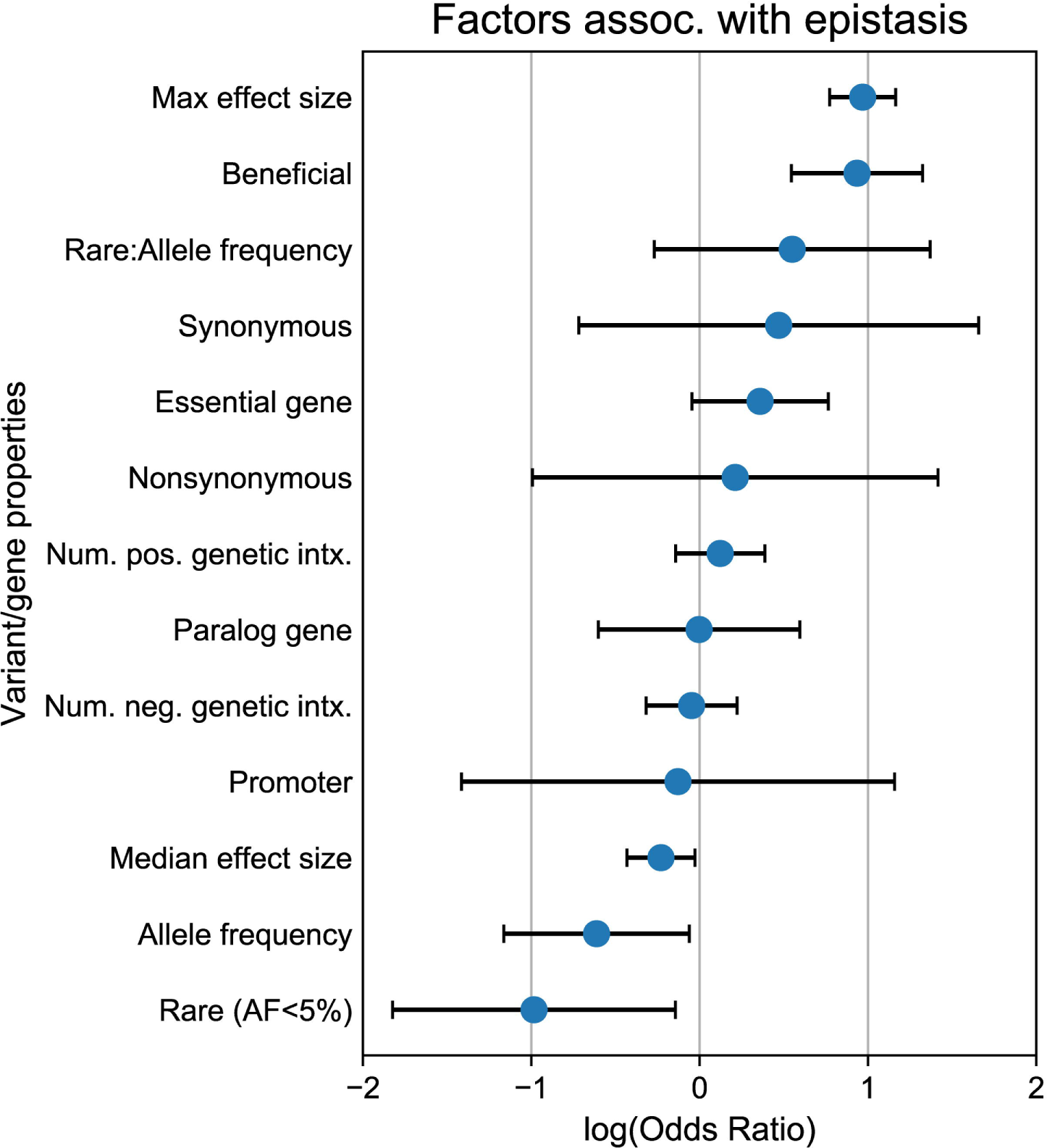
Odds ratios for multiple logistic regression with all variables tested for association with genetic interactions. Data is represented by mean log odds ratio ± 95% confidence interval from the logistic model.

**Figure S6.**
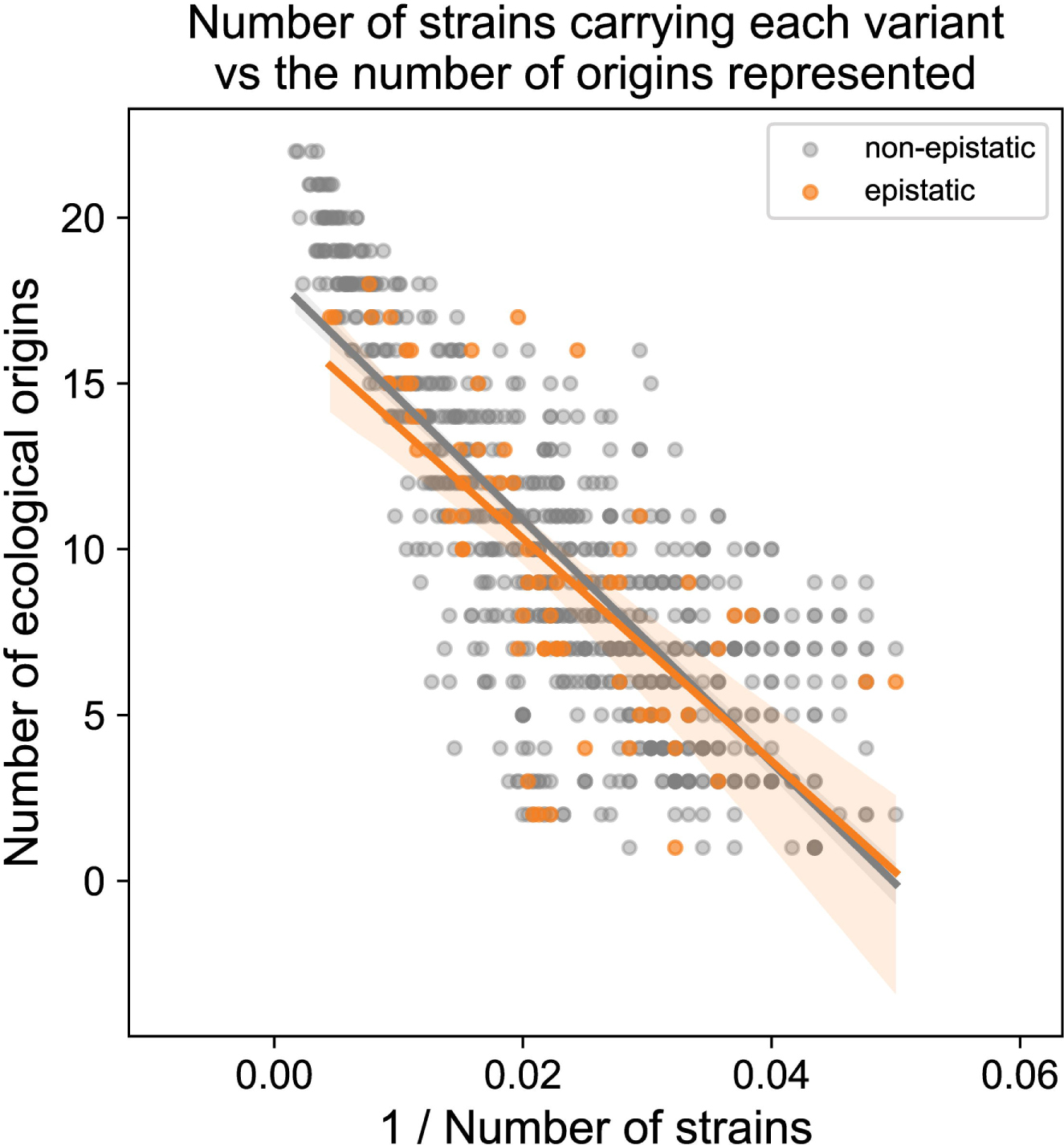
No significant difference in number of ecological origins represented between epistatic and non-epistatic variants. The number of strains carrying each variant is counted, as well as the number ecological origins in which the variant was observed in. Information of each yeast strain ecological origin was taken from Peter et al. (2018).

**Figure S7.**
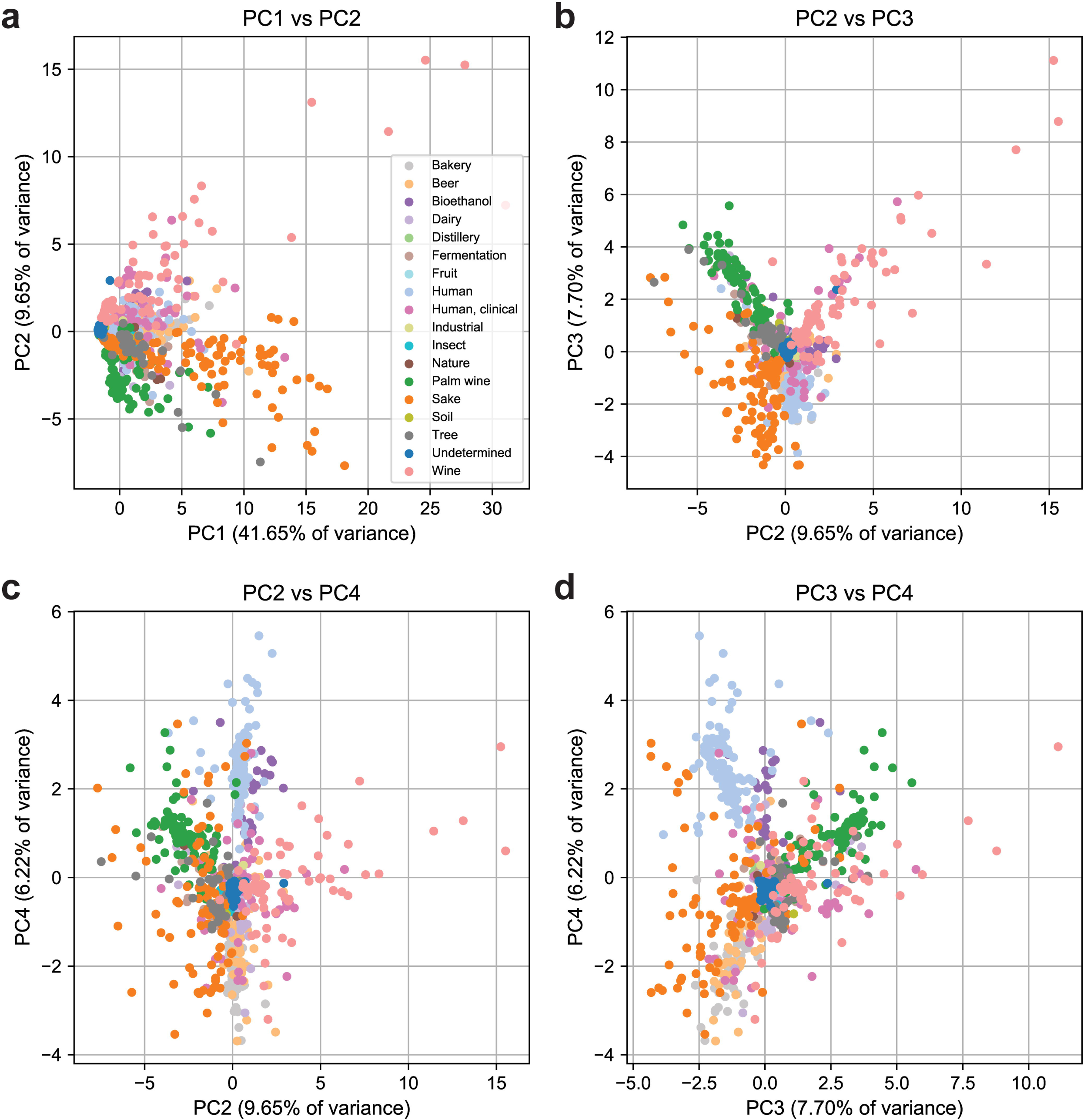
Principal component analysis of the population structure of natural variants. (a) PC1 separates variants by allele frequency, while (b-d) remaining PCs separate the variants based on their degree of representation within each ecological origin. Each point represents a variant, colored by the top ecological origin where the variant is present the most number of times.

**Figure S8.**
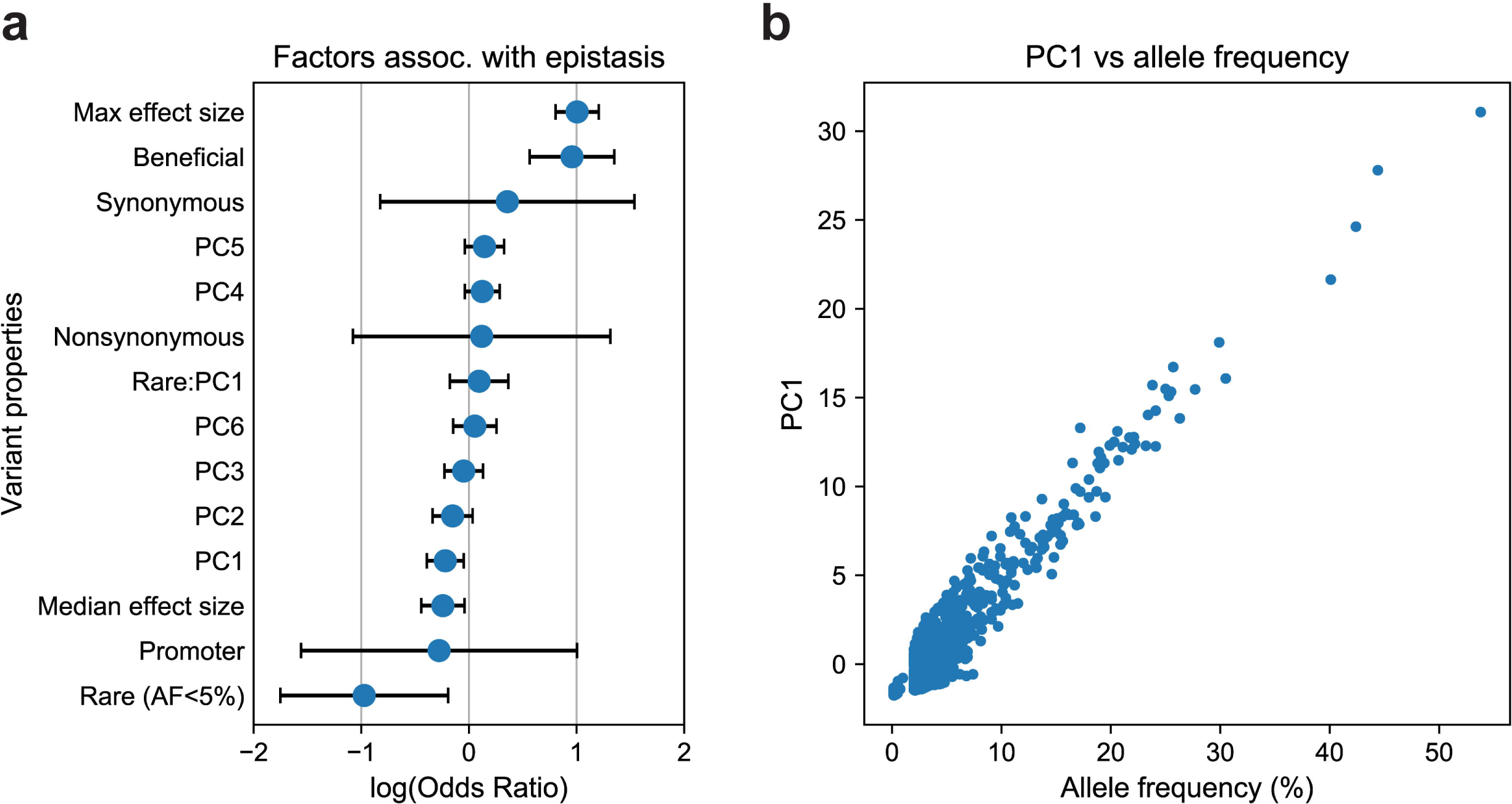
Allele frequency is the major determinant of the distribution of genetic interactions. (a) Multiple logistic regression odds ratio for variables associated with the incidence of genetic interactions, including top six principal components explaining ∼75% of variance in the variant distribution across ecological origins. PC1 is the only significant principal component in the model. Data is represented by mean log odds ratio ± 95% confidence interval from the logistic model. (b) PC1 is strongly correlated with allele frequency.

**Figure S9.**
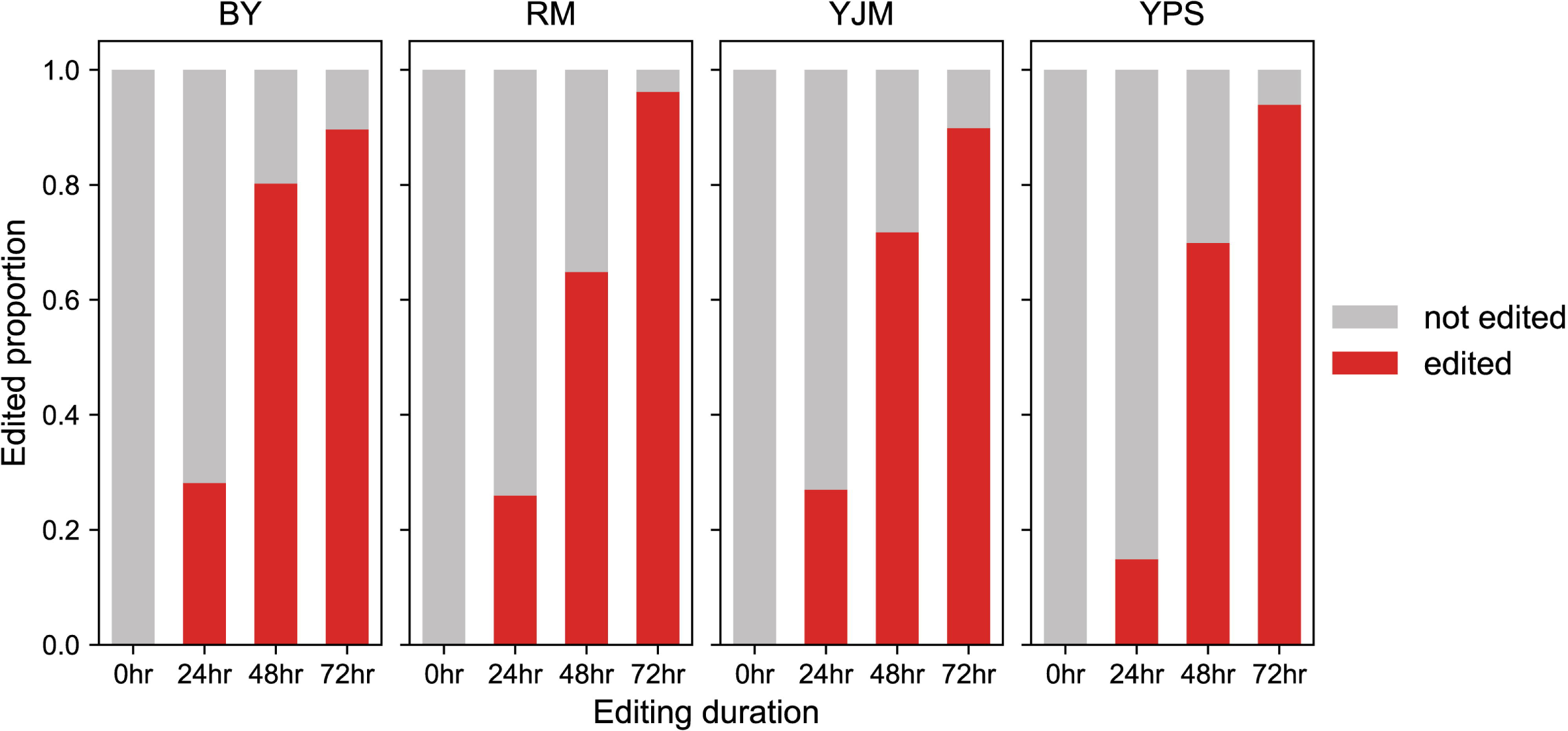
CRISPEY-BAR editing efficiency is equally high across the four yeast strains. All four strains achieve >90% editing rate at the end of 72 hours of induction. Editing efficiency was quantified from a CRISPEY-BAR plasmid editing a nonsense mutation into *ADE2*, with dilution plating every 24 hours to count pink colonies that indicate successful editing events.

## References

Ansanay Galeote, V., Alexandre, H., Bach, B., Delobel, P., Dequin, S., and Blondin, B. (2007). Sfl1p acts as an activator of the HSP30 gene in Saccharomyces cerevisiae. Current Genetics 52, 55–63. https://doi.org/10.1007/s00294-007-0136-z.

Atsushi, F., Yoshiko, K., Satoru, K., Yoshio, M., Shinichi, M., and Harumi, K. (1989). Domains of the SFL1 protein of yeasts are homologous to Myc oncoproteins or yeast heat-shock transcription factor. Gene 85, 321– 328. https://doi.org/10.1016/0378-1119(89)90424-1.

Babokhov, M., Mosaheb, M.M., Baker, R.W., and Fuchs, S.M. (2018). Repeat-Specific Functions for the C-Terminal Domain of RNA Polymerase II in Budding Yeast. G3 Genes|Genomes|Genetics *8*, 1593–1601. https://doi.org/10.1534/g3.118.200086.

Bakerlee, C.W., Nguyen Ba, A.N., Shulgina, Y., Rojas Echenique, J.I., and Desai, M.M. (2022). Idiosyncratic epistasis leads to global fitness–correlated trends. Science (1979) 376, 630–635. https://doi.org/10.1126/science.abm4774.

Bao, Z., HamediRad, M., Xue, P., Xiao, H., Tasan, I., Chao, R., Liang, J., and Zhao, H. (2018). Genome-scale engineering of Saccharomyces cerevisiae with single-nucleotide precision. Nature Biotechnology 36, 505–508. https://doi.org/10.1038/nbt.4132.

Benjamini, Y., and Hochberg, Y. (1995). Controlling the False Discovery Rate: A Practical and Powerful Approach to Multiple Testing. Journal of the Royal Statistical Society: Series B (Methodological) 57, 289–300. https://doi.org/10.1111/j.2517-6161.1995.tb02031.x.

Bergman, A., and Siegal, M.L. (2003). Evolutionary capacitance as a general feature of complex gene networks. Nature 424, 549–552. https://doi.org/10.1038/nature01765.

Bloom, J.S., Kotenko, I., Sadhu, M.J., Treusch, S., Albert, F.W., and Kruglyak, L. (2015). Genetic interactions contribute less than additive effects to quantitative trait variation in yeast. Nature Communications 6, 8712. https://doi.org/10.1038/ncomms9712.

Bony, M., Thines-Sempoux, D., Barre, P., and Blondin, B. (1997). Localization and cell surface anchoring of the Saccharomyces cerevisiae flocculation protein Flo1p. Journal of Bacteriology 179, 4929–4936. https://doi.org/10.1128/jb.179.15.4929-4936.1997.

Brem, R.B., Yvert, G., Clinton, R., and Kruglyak, L. (2002). Genetic Dissection of Transcriptional Regulation in Budding Yeast. Science (1979) 296, 752–755. https://doi.org/10.1126/science.1069516.

Breslow, D.K., Cameron, D.M., Collins, S.R., Schuldiner, M., Stewart-Ornstein, J., Newman, H.W., Braun, S., Madhani, H.D., Krogan, N.J., and Weissman, J.S. (2008). A comprehensive strategy enabling high-resolution functional analysis of the yeast genome. Nature Methods 5, 711–718. https://doi.org/10.1038/nmeth.1234.

Bystrykh, L. v. (2012). Generalized DNA Barcode Design Based on Hamming Codes. PLoS ONE 7, e36852. https://doi.org/10.1371/journal.pone.0036852.

Chandler, C.H., Chari, S., and Dworkin, I. (2013). Does your gene need a background check? How genetic background impacts the analysis of mutations, genes, and evolution. Trends in Genetics 29, 358–366. https://doi.org/10.1016/j.tig.2013.01.009.

Chen, R., Shi, L., Hakenberg, J., Naughton, B., Sklar, P., Zhang, J., Zhou, H., Tian, L., Prakash, O., Lemire, M., et al. (2016). Analysis of 589,306 genomes identifies individuals resilient to severe Mendelian childhood diseases. Nature Biotechnology 34, 531–538. https://doi.org/10.1038/nbt.3514.

Chou, H.-H., Chiu, H.-C., Delaney, N.F., Segrè, D., and Marx, C.J. (2011). Diminishing Returns Epistasis Among Beneficial Mutations Decelerates Adaptation. Science (1979) 332, 1190–1192. https://doi.org/10.1126/science.1203799.

Cooper, D.N., Krawczak, M., Polychronakos, C., Tyler-Smith, C., and Kehrer-Sawatzki, H. (2013). Where genotype is not predictive of phenotype: towards an understanding of the molecular basis of reduced penetrance in human inherited disease. Human Genetics 132, 1077–1130. https://doi.org/10.1007/s00439-013-1331-2.

Costanzo, M., VanderSluis, B., Koch, E.N., Baryshnikova, A., Pons, C., Tan, G., Wang, W., Usaj, M., Hanchard, J., Lee, S.D., et al. (2016). A global genetic interaction network maps a wiring diagram of cellular function. Science (1979) 353, aaf1420–aaf1420. https://doi.org/10.1126/science.aaf1420.

Doench, J.G., Fusi, N., Sullender, M., Hegde, M., Vaimberg, E.W., Donovan, K.F., Smith, I., Tothova, Z., Wilen, C., Orchard, R., et al. (2016). Optimized sgRNA design to maximize activity and minimize off-target effects of CRISPR-Cas9. Nature Biotechnology 34, 184–191. https://doi.org/10.1038/nbt.3437.

Dowell, R.D., Ryan, O., Jansen, A., Cheung, D., Agarwala, S., Danford, T., Bernstein, D.A., Alexander Rolfe, P., Heisler, L.E., Chin, B., et al. (2010). Genotype to phenotype: A Complex problem. Science (1979) 328, 469. https://doi.org/10.1126/science.1189015.

Ehrenreich, I.M. (2017). Epistasis: Searching for Interacting Genetic Variants Using Crosses. G3 Genes|Genomes|Genetics 7, 1619–1622. https://doi.org/10.1534/g3.117.042770.

Erwood, S., Bily, T.M.I., Lequyer, J., Yan, J., Gulati, N., Brewer, R.A., Zhou, L., Pelletier, L., Ivakine, E.A., and Cohn, R.D. (2022). Saturation variant interpretation using CRISPR prime editing. Nature Biotechnology https://doi.org/10.1038/s41587-021-01201-1.

Galardini, M., Busby, B.P., Vieitez, C., Dunham, A.S., Typas, A., and Beltrao, P. (2019). The impact of the genetic background on gene deletion phenotypes in Saccharomyces cerevisiae. Molecular Systems Biology 15. https://doi.org/10.15252/msb.20198831.

Gietz, R.D., and Schiestl, R.H. (2007). High-efficiency yeast transformation using the LiAc/SS carrier DNA/PEG method. Nature Protocols 2, 31–34. https://doi.org/10.1038/nprot.2007.13.

Goldstein, I., and Ehrenreich, I.M. (2021). The complex role of genetic background in shaping the effects of spontaneous and induced mutations. Yeast 38, 187–196. https://doi.org/10.1002/yea.3530.

Guo, X., Chavez, A., Tung, A., Chan, Y., Kaas, C., Yin, Y., Cecchi, R., Garnier, S.L., Kelsic, E.D., Schubert, M., et al. (2018). High-throughput creation and functional profiling of DNA sequence variant libraries using CRISPR–Cas9 in yeast. Nature Biotechnology 36, 540–546. https://doi.org/10.1038/nbt.4147.

Hanna, R.E., Hegde, M., Fagre, C.R., DeWeirdt, P.C., Sangree, A.K., Szegletes, Z., Griffith, A., Feeley, M.N., Sanson, K.R., Baidi, Y., et al. (2021). Massively parallel assessment of human variants with base editor screens. Cell 184, 1064–1080.e20. https://doi.org/10.1016/j.cell.2021.01.012.

Hemani, G., Knott, S., and Haley, C. (2013). An Evolutionary Perspective on Epistasis and the Missing Heritability. PLoS Genetics 9, e1003295. https://doi.org/10.1371/journal.pgen.1003295.

Huang, W., Richards, S., Carbone, M.A., Zhu, D., Anholt, R.R.H., Ayroles, J.F., Duncan, L., Jordan, K.W., Lawrence, F., Magwire, M.M., et al. (2012). Epistasis dominates the genetic architecture of *Drosophila* quantitative traits. Proceedings of the National Academy of Sciences 109, 15553–15559. https://doi.org/10.1073/pnas.1213423109.

Johnson, M.S., Martsul, A., Kryazhimskiy, S., and Desai, M.M. (2019). Higher-fitness yeast genotypes are less robust to deleterious mutations. Science (1979) 366, 490–493. https://doi.org/10.1126/science.aay4199.

Kao, K.C., and Sherlock, G. (2008). Molecular characterization of clonal interference during adaptive evolution in asexual populations of Saccharomyces cerevisiae. Nature Genetics 40, 1499–1504. https://doi.org/10.1038/ng.280.

Kryazhimskiy, S., Rice, D.P., Jerison, E.R., and Desai, M.M. (2014). Global epistasis makes adaptation predictable despite sequence-level stochasticity. Science (1979) 344, 1519–1522. https://doi.org/10.1126/science.1250939.

Kuzmin, E., VanderSluis, B., Wang, W., Tan, G., Deshpande, R., Chen, Y., Usaj, M., Balint, A., Mattiazzi Usaj, M., van Leeuwen, J., et al. (2018). Systematic analysis of complex genetic interactions. Science (1979) 360. https://doi.org/10.1126/science.aao1729.

Langmead, B., and Salzberg, S.L. (2012). Fast gapped-read alignment with Bowtie 2. Nature Methods 9, 357– 359. https://doi.org/10.1038/nmeth.1923.

Levy, S.F., Blundell, J.R., Venkataram, S., Petrov, D.A., Fisher, D.S., and Sherlock, G. (2015). Quantitative evolutionary dynamics using high-resolution lineage tracking. Nature 519, 181–186. https://doi.org/10.1038/nature14279.

Li, C., Qian, W., Maclean, C.J., and Zhang, J. (2016). The fitness landscape of a tRNA gene. Science (1979) 352, 837–840. https://doi.org/10.1126/science.aae0568.

Liu, H., Styles, C.A., and Fink, G.R. (1996). Saccharomyces cerevisiae S288C has a mutation in FLO8, a gene required for filamentous growth. Genetics 144, 967–978. https://doi.org/10.1093/genetics/144.3.967.

Love, M.I., Huber, W., and Anders, S. (2014). Moderated estimation of fold change and dispersion for RNA-seq data with DESeq2. Genome Biology 15, 550. https://doi.org/10.1186/s13059-014-0550-8.

Lyons, D.M., Zou, Z., Xu, H., and Zhang, J. (2020). Idiosyncratic epistasis creates universals in mutational effects and evolutionary trajectories. Nature Ecology & Evolution 4, 1685–1693. https://doi.org/10.1038/s41559-020-01286-y.

Magoc, T., and Salzberg, S.L. (2011). FLASH: fast length adjustment of short reads to improve genome assemblies. Bioinformatics 27, 2957–2963. https://doi.org/10.1093/bioinformatics/btr507.

Martin, M. (2011). Cutadapt removes adapter sequences from high-throughput sequencing reads. EMBnet J 17, 10. https://doi.org/10.14806/ej.17.1.200.

Masel, J. (2013). Q&A: Evolutionary capacitance. BMC Biology 11, 103. https://doi.org/10.1186/1741-7007-11-103.

Nadeau, J.H. (2001). Modifier genes in mice and humans. Nature Reviews Genetics 2, 165–174. https://doi.org/10.1038/35056009.

Narasimhan, V.M., Hunt, K.A., Mason, D., Baker, C.L., Karczewski, K.J., Barnes, M.R., Barnett, A.H., Bates, C., Bellary, S., Bockett, N.A., et al. (2016). Health and population effects of rare gene knockouts in adult humans with related parents. Science (1979) *352*, 474–477. https://doi.org/10.1126/science.aac8624.

Peter, J., de Chiara, M., Friedrich, A., Yue, J.X., Pflieger, D., Bergström, A., Sigwalt, A., Barre, B., Freel, K., Llored, A., et al. (2018). Genome evolution across 1,011 Saccharomyces cerevisiae isolates. Nature 556, 339– 344. https://doi.org/10.1038/s41586-018-0030-5.

Pothoulakis, G., and Ellis, T. (2018). Construction of hybrid regulated mother-specific yeast promoters for inducible differential gene expression. PLoS ONE 13, e0194588. https://doi.org/10.1371/journal.pone.0194588.

Puchta, O., Cseke, B., Czaja, H., Tollervey, D., Sanguinetti, G., and Kudla, G. (2016). Network of epistatic interactions within a yeast snoRNA. Science (1979) 352, 840–844. https://doi.org/10.1126/science.aaf0965.

Reddy, G., and Desai, M.M. (2021). Global epistasis emerges from a generic model of a complex trait. Elife 10. https://doi.org/10.7554/eLife.64740.

Richardson, C.D., Ray, G.J., DeWitt, M.A., Curie, G.L., and Corn, J.E. (2016). Enhancing homology-directed genome editing by catalytically active and inactive CRISPR-Cas9 using asymmetric donor DNA. Nature Biotechnology 34, 339–344. https://doi.org/10.1038/nbt.3481.

Riordan, J.D., and Nadeau, J.H. (2017). From Peas to Disease: Modifier Genes, Network Resilience, and the Genetics of Health. The American Journal of Human Genetics 101, 177–191. https://doi.org/10.1016/j.ajhg.2017.06.004.

Robertson, L.S., and Fink, G.R. (1998). The three yeast A kinases have specific signaling functions in pseudohyphal growth. Proceedings of the National Academy of Sciences 95, 13783–13787. https://doi.org/10.1073/pnas.95.23.13783.

Rockman, M. V. (2012). THE QTN PROGRAM AND THE ALLELES THAT MATTER FOR EVOLUTION: ALL THAT’S GOLD DOES NOT GLITTER. Evolution (N Y) 66, 1–17. https://doi.org/10.1111/j.1558-5646.2011.01486.x.

Roosen, J., Engelen, K., Marchal, K., Mathys, J., Griffioen, G., Cameroni, E., Thevelein, J.M., de Virgilio, C., de Moor, B., and Winderickx, J. (2004). PKA and Sch9 control a molecular switch important for the proper adaptation to nutrient availability. Molecular Microbiology 55, 862–880. https://doi.org/10.1111/j.1365-2958.2004.04429.x.

Rutherford, S.L., and Lindquist, S. (1998). Hsp90 as a capacitor for morphological evolution. Nature 396, 336– 342. https://doi.org/10.1038/24550.

Schaid, D.J., Chen, W., and Larson, N.B. (2018). From genome-wide associations to candidate causal variants by statistical fine-mapping. Nature Reviews Genetics 19, 491–504. https://doi.org/10.1038/s41576-018-0016-z.

Sharon, E., Chen, S.A.A., Khosla, N.M., Smith, J.D., Pritchard, J.K., and Fraser, H.B. (2018). Functional Genetic Variants Revealed by Massively Parallel Precise Genome Editing. Cell 175, 544–557.e16. https://doi.org/10.1016/j.cell.2018.08.057.

Sniegowski, P.D., Dombrowski, P.G., and Fingerman, E. (2002). *Saccharomyces cerevisiae* and *Saccharomyces paradoxus* coexist in a natural woodland site in North America and display different levels of reproductive isolation from European conspecifics. FEMS Yeast Research 1, 299–306. https://doi.org/10.1111/j.1567-1364.2002.tb00048.x.

Soares, E.V. (2011). Flocculation in Saccharomyces cerevisiae: a review. Journal of Applied Microbiology 110, 1–18. https://doi.org/10.1111/j.1365-2672.2010.04897.x.

Song, W., and Carlson, M. (1998). Srb/mediator proteins interact functionally and physically with transcriptional repressor Sfl1. The EMBO Journal 17, 5757–5765. https://doi.org/10.1093/emboj/17.19.5757.

Su, S.S., and Mitchell, A.P. (1993). Identification of functionally related genes that stimulate early meiotic gene expression in yeast. Genetics 133, 67–77. https://doi.org/10.1093/genetics/133.1.67.

Tawfik, O.W., Papasian, C.J., Dixon, A.Y., and Potter, L.M. (1989). Saccharomyces cerevisiae pneumonia in a patient with acquired immune deficiency syndrome. Journal of Clinical Microbiology 27, 1689–1691. https://doi.org/10.1128/jcm.27.7.1689-1691.1989.

Taylor, M.B., and Ehrenreich, I.M. (2015). Higher-order genetic interactions and their contribution to complex traits. Trends in Genetics 31, 34–40. https://doi.org/10.1016/j.tig.2014.09.001.

Vidan, S., and Mitchell, A.P. (1997). Stimulation of yeast meiotic gene expression by the glucose-repressible protein kinase Rim15p. Molecular and Cellular Biology 17, 2688–2697. https://doi.org/10.1128/MCB.17.5.2688.

Wei, X., and Zhang, J. (2019). Patterns and Mechanisms of Diminishing Returns from Beneficial Mutations. Molecular Biology and Evolution 36, 1008–1021. https://doi.org/10.1093/molbev/msz035.

Wei, W., McCusker, J.H., Hyman, R.W., Jones, T., Ning, Y., Cao, Z., Gu, Z., Bruno, D., Miranda, M., Nguyen, M., et al. (2007). Genome sequencing and comparative analysis of *Saccharomyces cerevisiae* strain YJM789. Proceedings of the National Academy of Sciences 104, 12825–12830. https://doi.org/10.1073/pnas.0701291104.

Weinreich, D.M., Watson, R.A., and Chao, L. (2005). Perspective: Sign epistasis and genetic constraint on evolutionary trajectories. Evolution 59, 1165–1174. .

Weinreich, D.M., Delaney, N.F., DePristo, M.A., and Hartl, D.L. (2006). Darwinian Evolution Can Follow Only Very Few Mutational Paths to Fitter Proteins. Science (1979) 312, 111–114. https://doi.org/10.1126/science.1123539.

Xie, Z.-X., Mitchell, L.A., Liu, H.-M., Li, B.-Z., Liu, D., Agmon, N., Wu, Y., Li, X., Zhou, X., Li, B., et al. (2018). Rapid and Efficient CRISPR/Cas9-Based Mating-Type Switching of *Saccharomyces cerevisiae*. G3 Genes|Genomes|Genetics 8, 173–183. https://doi.org/10.1534/g3.117.300347.

Yue, J.-X., Li, J., Aigrain, L., Hallin, J., Persson, K., Oliver, K., Bergström, A., Coupland, P., Warringer, J., Lagomarsino, M.C., et al. (2017). Contrasting evolutionary genome dynamics between domesticated and wild yeasts. Nature Genetics 49, 913–924. https://doi.org/10.1038/ng.3847.

